# Impact of emerging mutations on the dynamic properties the SARS-CoV-2 main protease: an *in silico* investigation

**DOI:** 10.1101/2020.05.29.123190

**Authors:** Olivier Sheik Amamuddy, Gennady M. Verkhivker, Özlem Tastan Bishop

**Affiliations:** Research Unit in Bioinformatics, Department of Microbiology and Biochemistry, Rhodes University, Grahamstown, South Africa; Graduate Program in Computational and Data Sciences, Schmid College of Science and Technology, Chapman University, Orange, CA 92866, United States of America; Department of Biomedical and Pharmaceutical Sciences, Chapman University School of Pharmacy, Irvine, CA 92618, United States of America; Department of Pharmacology, Skaggs School of Pharmacy and Pharmaceutical Sciences, University of California San Diego, 9500 Gilman Drive, La Jolla, CA 92093, USA

**Keywords:** SARS-CoV-2, main protease, molecular dynamics, non-synonymous mutations, MD-TASK

## Abstract

The new coronavirus (SARS-CoV-2) is a global threat to world health and its economy. Its main protease (M^pro^), which functions as a dimer, cleaves viral precursor proteins in the process of viral maturation. It is a good candidate for drug development owing to its conservation and the absence of a human homolog. An improved understanding of the protein behaviour can accelerate the discovery of effective therapies in order to reduce mortality. 100 ns all-atom molecular dynamics simulations of 50 homology modelled mutant M^pro^ dimers were performed at pH 7 from filtered sequences obtained from the GISAID database. Protease dynamics were analysed using RMSD, RMSF, R_g_, the averaged *betweenness centrality* and geometry calculations. Domains from each M^pro^ protomer were found to generally have independent motions, while the dimer-stabilising N-finger region was found to be flexible in most mutants. A mirrored interprotomer pocket was found to be correlated to the catalytic site using compaction dynamics, and can be a potential allosteric target. The high number of titratable amino acids of M^pro^ may indicate an important role of pH on enzyme dynamics, as previously reported for SARS-CoV. Independent coarse-grained Monte Carlo simulations suggest a link between rigidity/mutability and enzymatic function.

## 1. Introduction

The human severe acute respiratory syndrome coronavirus 2 (SARS-CoV-2) strain is the causative agent of the COVID-19 pandemic [1]. After being first reported from the Wuhan seafood and animal market in late December 2019 [2], the total number of reported cases worldwide has reached over 5.8 million, with an overall crude death rate reported at 3.67% [3]. The disease is however more severe amongst the elderly, and those living with co-morbidities that involve endothelial dysfunction [4], such as hypertension, obesity and diabetes [5]. While there is currently no cure or available vaccine [2,6], there is a lot of uncertainty around the behaviour of the pathogen. Drastic measures designed to limit the rate of new infections [7] have resulted in global economic problems, which have affected many livelihoods, even exacerbating food insecurity [8]. Owing to the rapid generation of genomic sequence data [9,10] and the timely availability of 3D structural data, research into potential drugs is under way alongside clinical trials. Fundamental research is key to understanding the pathogen’s strategies such that more informed decisions can be made about clinical interventions. As seen in other pathogens, mutations occur through the normal process of evolution, and certain advantageous variations can be selected for over time. The SARS-CoV-2 genome is RNA-based, and viruses from this category have been reported to have increased rates of mutation [11]. For instance, in HIV this has lead to several levels of classification of the virus, in which certain strains can manifest different transmissibility patterns and show differing responses to existing therapies [12,13]. From the data gathered from the GISAID database [9] and real-time sub-sample estimates of genetic relatedness from the Nextstrain web resource [14], it is clear that the virus is evolving within the human host.

Although progress is being gradually made in understanding viral structural biology and symptomatology of the disease, current knowledge is still fragmentary [15,16], while the death toll and the number of infections keeps on rising. Thus, time is of the essence for the discovery of effective therapies. It is imperative to better characterise parts of the viral mechanisms to better understand the behaviour of the new coronavirus. Already, with the help of experimentally determined structures, genomic data and annotations, a growing number of *in silico* work is suggesting potential solutions to the COVID-19 pandemic using various techniques, including the use of molecular modelling, network-analysis [17–20]and machine learning [2,21–24]. Collectively, these may pave the way to a potential solution. In this study we report on some of the recently emerged mutations of the SARS-CoV-2 M^pro^ protein, and investigate different aspects of their dynamics of the using molecular dynamics.

The M^pro^ enzyme, also known as the 3C-like protease, is one of the best studied drug targets among the coronaviruses [25]. This is mainly due to the similarities in active site and mechanisms with the related pathogenic betacoronaviruses from previous epidemics of SARS-CoV and MERS-CoV (Middle East respiratory syndrome coronavirus) [26]. M^pro^ is a conserved drug target present in all members of the *Coronavirinae* subfamily [27,28] and is highly similar to its SARS-CoV counterpart [26]. SARS-CoV-2 M^pro^ does not have a human homolog [20], which reduces the chances of accidentally targeting host proteins. Alongside the papain-like protease (PLP) enzyme, M^pro^ plays an essential role in the process of viral maturation [2], cleaving the large precursor replicase polyprotein 1ab to produce 16 non-structural proteins [2,29]. The cysteine protease functions as a homodimer and mainly comprises three domains (I-III) [2]. Homo-dimerisation plays an important role in the catalytic activity of M^pro^, as reported in the case of the SARS CoV M^pro^ homolog, where the G11A mutation completely abolished its activity by interfering with the insertion of the “N-finger” region (residues 1-9) [30]. At the N-terminus the chymotrypsin-like domain I (residues 10-99) is connected to the picornavirus 3C-protease like domain II (residues 100-182), which together form a hydrophobic substrate binding site, with catalytic residues CYS145 and HIS41 [29,31]. Domain III (residues 198-303; also referred to as the helical domain) is connected to domain II [32] by a 15 residue linker loop. While each domain minimally contacts its equivalent domain from the alternate chain, the majority of the dimer contact interface is a result of interactions present between domain II (chain A) and the N-finger (chain B) [29]. In the same manner, the N-finger from chain A contacts domain II from chain B. Each chain is referred to as a protomer [22,29,33] 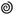.

In this work we study the collective effects of various M^pro^ mutations of from a filtered sample 50 isolates of SARS-CoV-2 by first mapping them on 3D structures and performing all-atom molecular dynamics (MD) for each of the mutants, in addition to the reference protein. All-atom simulations were carried out at a constant protonation state corresponding to a pH of 7. Multiple aspects of the protein dynamics were analysed using a battery of techniques, including the averaged *betweenness centrality (BC) -* a metric of Dynamic Residue Network (DRN) analysis, dynamic cross-correlation (DCC) [34], geometry calculations (inter-domain angles and interprotomer distances) based on the centre of mass (COM), cavity compaction analyses, and the analysis of residue and backbone fluctuations. Coarse-grained Monte Carlo simulations were also independently investigated.

## 2. Results and discussion

### 2.1. Analysis of residue mutations and their distribution in the 3D structure

As a preliminary investigation of the propensity of the sequences to acquire novel mutations, unique residue mutations were determined across our set of 50 protein sequences, which were filtered from the GISAID database [9]. While we cannot infer population frequencies (across the world) from our relatively small sample, we show from our estimate that multiple non-synonymous mutations have already occurred on each domain of the M^pro^ (Fig. 1). These include the following mutations: A7V, G15D/S, M17I, V20L, T45I, D48E, M49I, R60C, K61R, A70T, G71S, L89F, K90R, P99L, Y101C, R105H, P108S, A116V, A129V, P132L, T135I, I136V, N151D, V157I/L, C160S, A173V, P184L/S, T190I, A191V, A193V, T196M, T198I, T201A, L220F, L232F, A234V, K236R, Y237H, D248E, A255V, T259I, A260V, V261A, A266V, N274D, R279C and S301L. From these, it can be observed that many mutations are inter-conversions of the hydrophobic side-chain residues alanine and valine. As seen in Fig. 1, most residue mutations have occurred in solvent-accessible surfaces, with the exception of A7V, V20L, L89F, A116V, A129V, T135I, I136V, V157I/L, C160S, A173V, T201A, A234V and A266V, which were predicted to be buried by the PyMOL script *findSurfaceResidues*, using the default cut-off of 2.5 Å^2^.

**Fig. 1.**
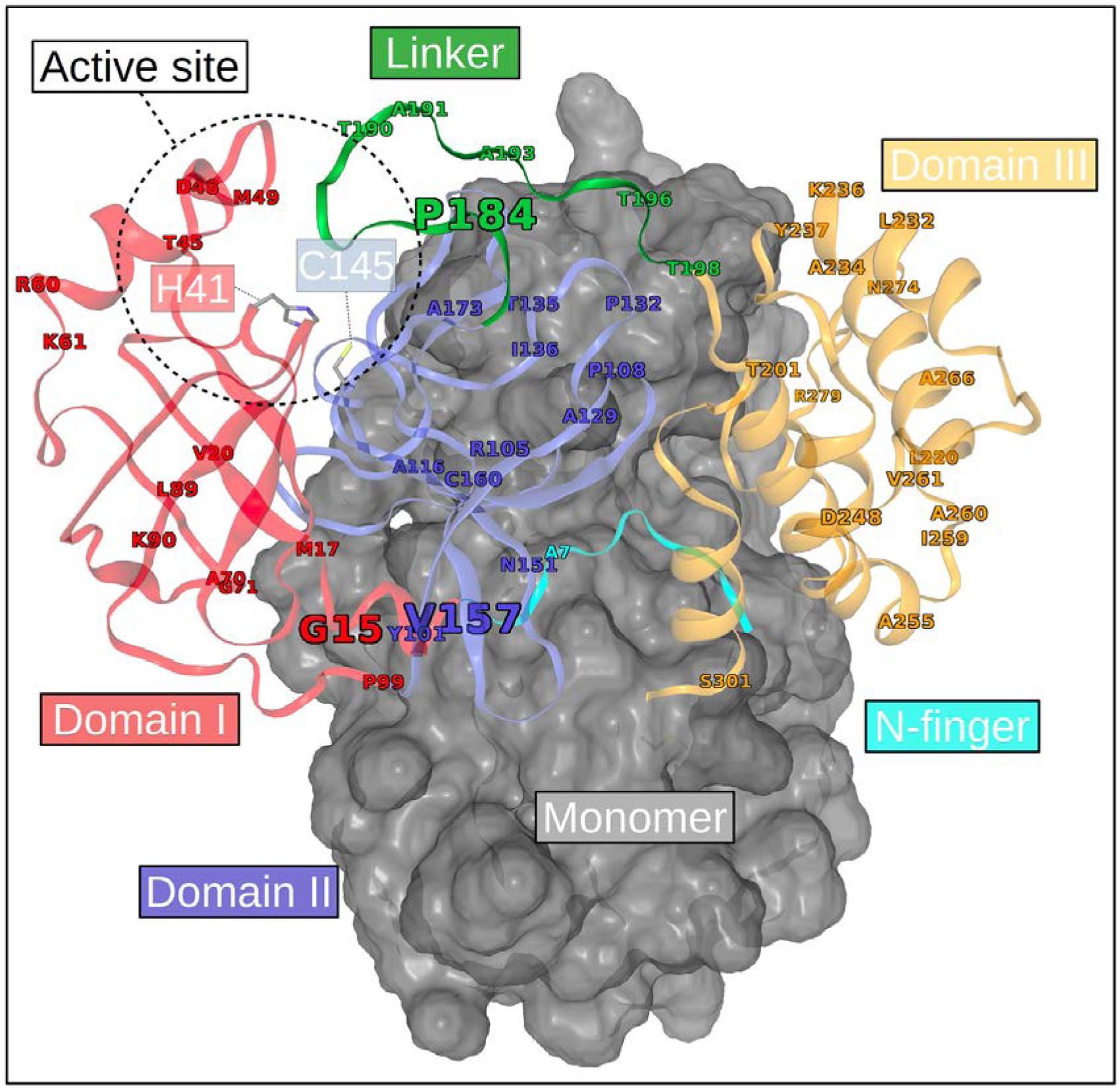
Mapping of the positions showing unique mutations from the reference M^pro^ sequence. For clarity, domains (I-III) are coloured (red, blue and orange respectively) only for one of the monomers, while the other is represented as a grey surface. The domain linker region is in green and the N-finger is in cyan. The size of the labels denotes the number of unique mutations recorded at that position.

Further, a higher rate of non-synonymous mutations has occurred at residue position 15 (G15D/S) in domain I, residue position 157 (V157I/L) in domain II and at position 184 (P184L/S) within the inter-domain linker region. On the 3D structure, it can be seen that these mutable areas occur away from the core areas of domains I and II, hence the probable lower selective pressure for these regions. For this reason, we posit that individually, these loci may be less important for basic enzymatic function. Mutation rates of RNA viruses are known to be generally high, endowing them with the ability to escape host immune responses, improve their virulence, and even change tissue tropism [11,35]. However, extinction events are not uncommon among the RNA viruses, as seen in the influenza A H1N1 strains [36] and the previous SARS-CoV strain [37], and are presumed to be associated with the gradual accumulation of non-synonymous mutations. On the other hand, a reduction in replicative speed and genetic diversity was independently observed in the poliovirus 3D^G64S^ mutant compared to its wild-type (WT) when rates of mutation were artificially increased by exposure to a mutagen [11,38,39]. In the case of the HIV, multiple mutations of minor effect are known to collaboratively modulate the effect of other advantageous mutations already present in the protease [40,41]. As a newly emerged pathogen with a relatively long incubation period and an incompletely understood biology, these facts from related viruses presuppose a potentially complex mechanism of evolution and adaptation, which suggests that mutations have to be closely monitored for global health and security.

It is interesting to note that amongst the buried residue mutations, domain II has accumulated the highest number of these mutations in such little time, which may suggest a certain degree of tolerance to mutations in that region, despite their presence within beta strands. In the same domain, the A116V mutation occurs on a beta strand, which is supported by a rich network of hydrogen bonds. The local impacts of the A116V mutation are discussed further in section 2.2.

While several mutations are present in domain III, the current data suggests that these have not (yet) further evolved at these positions, and probably suggests that there might a higher fitness cost involved with mutating residues from this domain. It is also possible that under-sampling may lead to a similar observation. Most of the mutations have occurred in solvent-exposed surfaces of the protein, which may be under reduced selective pressure, especially if these are in loop regions. On the other hand, interfacial residue mutations (particularly at the chain interface) may exert their effects in a more exacerbated manner by either over-stabilising or destabilising protein-protein interactions, as reported in work on other proteins [42,43]. One such mutation has already happened at position 7 (A7V) in the N-finger region – a region that is vital for enzymatic activity [31]. The local impacts of the A7V mutation are discussed in section 2.2.

### 2.2. Homology modelling and inspection of residue protonation states

After the preliminary analysis of M^pro^ at the sequence level, their 3D structures were built for further investigation. The z-DOPE scores for the best homology models obtained for each sample were all below -1, indicating that they were all native-like [51,52]. Overall, z-DOPE values had a minimum of -1.48, a maximum of -1.38, with a median of -1.42. While it is not possible to simulate changes in protonation state using classical MD, an initial approximation of the most prevalent residue protonation states (at pH 7) was used for each of the M^pro^ samples. As there was a relatively higher level of variation in residue protonation states amongst each of the seven histidine residues found in each M^pro^ protomer, only the catalytic residue and the non-synonymous mutations are described herein, post homology modelling. We suspect that the high number of titratable amino acids may play a role in influencing protein behaviour at varying pH levels. Previous work in SARS CoV reports of a “pH-dependent activity-switch” of the main protease [53]. In our case, the catalytic residue HIS41 was generally protonated at the delta nitrogen (HID) atom, but also occurred in its fully protonated state (HIP) in one of the protomers for samples EPI_ISL_419710 and EPI_ISL_425655. Protonated aspartic acid (ASH) was found in both protomers of sample EPI_ISL_420510, which was the only isolate to contain the N151D mutation. ASH was also found in sample EPI_ISL_421312, for only one of its protomers at residue position 289, which otherwise occurs in its deprotonated form in all other samples. The R105H mutation (present only in sample EPI_ISL_419984) occurs as a HID in each protomer. Similarly the Y237H mutation, present only in sample EPI_ISL_416720 occurs as HID in both of its protomers.

The local residue interactions around residue mutations of interest were also investigated, by comparing them with their equivalent position in the M^pro^ reference, using their modelled structure. These mutations comprised residues that underwent a higher number of mutations (G15D/S, V157I/L and P184L/S) in addition to mutations that occurred at or close to the dimer interface (A7V and A116V). A7V significantly increased the number of proximal interactions to neighbouring residues when compared to the reference protein (Fig. 2), and gaining in hydrophobic interactions, although a clash in van der Waal radius is additionally present. G15D is found to increase the number of proximal contacts by the largest extent, while also modestly increasing the number of hydrophobic interactions. By replacing alanine with valine at position 116, increased amount of proximal interactions is gained at V116 in sample EPI_ISL_425284, with an increased amount of hydrophobic contacts to the residue. Mutations V157I/L both reduced the number of local hydrophobic contacts and resulted in a reduced van der Waal clash compared to the reference. P184L had a reduced number of proximal contacts compared to the reference, while P184S was very similar to its equivalent position in the reference. These observations only give a general indication of the local changes present in the static starting structures. In the sections that follow, the dynamic aspects of M^pro^ are investigated.

**Fig. 2.**
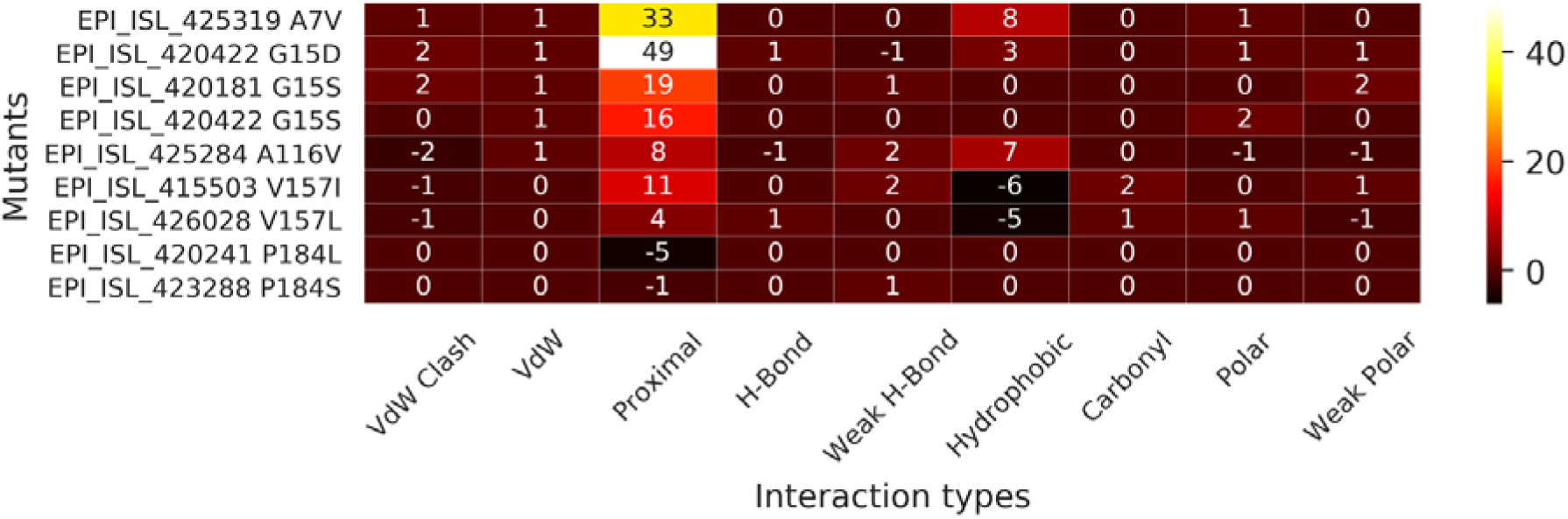
Differences in the sum of each interaction type for the M^pro^ only for the residue locations that either accumulated more than one non-synonymous mutation or ones occurring at protein interfaces. The differences were obtained by subtracting the reference values from the matching residue loci in the mutant. Sample names and the selected residue mutations are shown along the y-axis.

### 2.3. Estimation of the protein backbone flexibility from MD using C_α_ RMSD

The Cα RMSD obtained after frame fitting to the initial frames and periodic image correction (Fig. 3A) showed noticeably higher backbone flexibility for the isolates EPI_ISL_416720, EPI_ISL_420610, EPI_ISL_421380, EPI_ISL_421763, EPI_ISL_423772, EPI_ISL_425284, EPI_ISL_425319 EPI_ISL_425886 and EPI_ISL_426097. Additionally, from the shapes of the kernel density plots, it can be seen that some mutants may be equilibrating around single energy minima (for uni-modal distributions), while others are selecting for multiple major conformations, as seen from the multi-modal shapes. In all, these samples involve mutations A7V, M17I, A70T, A116V, K236R, Y237H, D248E, A266V and N274D, in which the last five mutations are exclusive to domain III. A7V occurs on the N-finger, which is a critical region for Mpro dimer stability. M17I occurs on an internal loop that connects a beta strand to a helix in domain I, while A70T occurs on solvent-exposed loop in the same domain. Mutation A116V occurs in a buried beta strand within domain II. From their visibly positively shifted upper quartiles, we may infer that more mobile backbones were sampled for mutants EPI_ISL_421380, EPI_ISL_423772 and EPI_ISL_425886. Upon visualising the trajectories, a slight twisting motion was observed between their protomers. More exactly, the protomers generally moved in opposite directions, whilst being tethered at the center. However, a similar motion was also observed in all other samples. This may indicate that the twisting motions are a normal behaviour of dimeric M^pro^, at least under our simulated conditions for the apo state. We suspect that the changes may rather be observable at more local levels, such at the intra/inter-domain or residue levels. This twisting motion is summarised using the first non-trivial mode (number 7) obtained from the anisotropic network model (ANM) of the reference protease (Fig. 3A). The predictions from ProDy [44] were visualised in VMD [45] 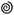.

**Fig. 3.**
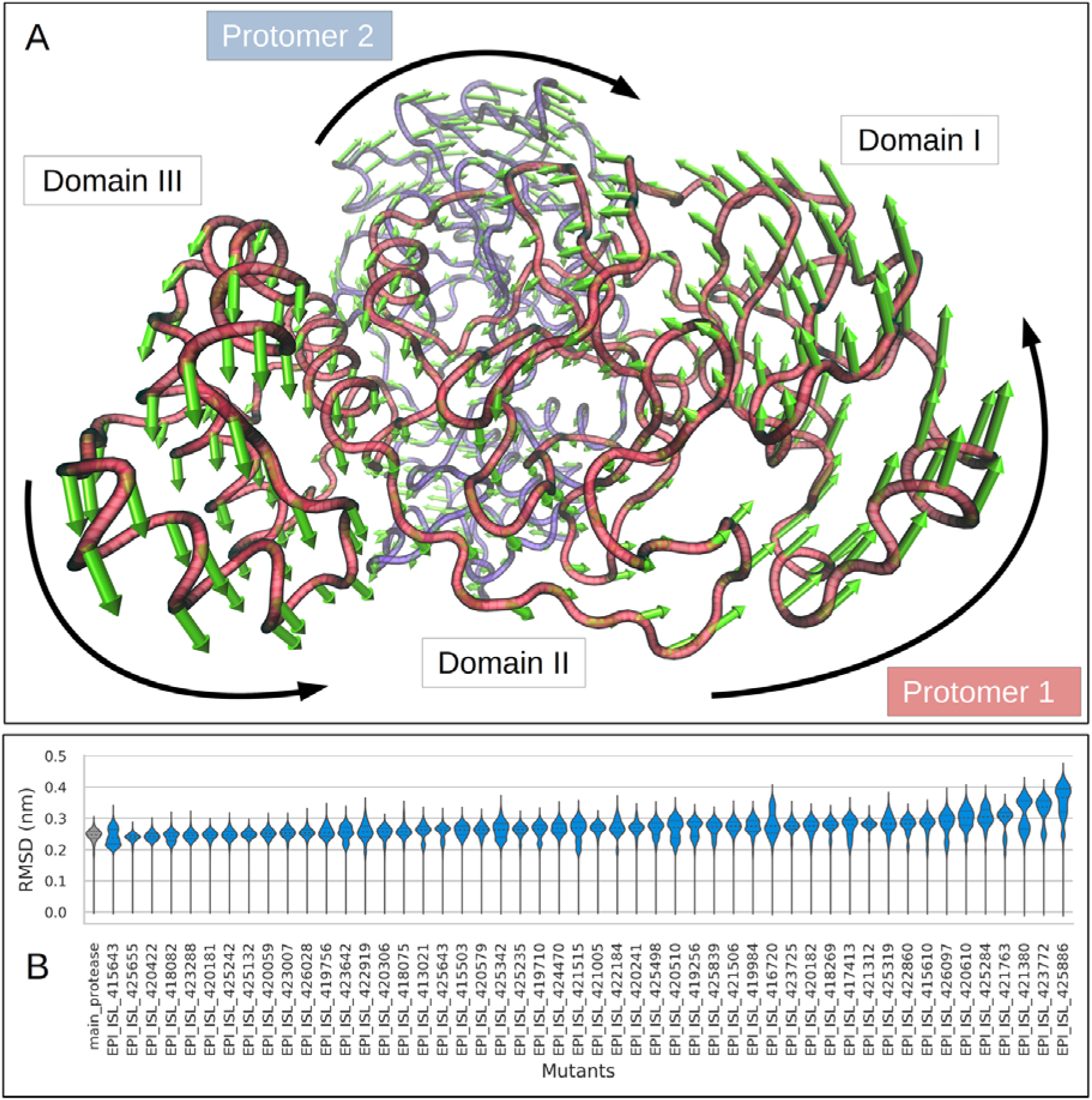
(A) The general twisting motion of the protomers observed across Mpro samples, inferred from the reference protease using ANM. (B) Violin plots of Cα RMSD values for the reference (in grey) and the mutant (coloured in blue) Mpro, showing the 25th, 50th and 75th percentiles in dotted lines inside the kernel density plots. Distributions are scaled by counts, and have been sorted by median RMSD for the mutants.

### 2.4. Estimation of the N-finger flexibility from MD using all-atom RMSD

The N-finger region is an important structural foundation required for the stabilisation of the functional M^pro^ dimer. Therefore we investigated its mobility within the dimeric protein across all M^pro^ samples. After fitting the proteins globally, no further fitting was done for the N-finger in order to better represent the N-finger motions. As seen in Fig 4., the reference protein has a very stable N-finger conformation, although the protomer equilibria are different. This tendency towards asymmetry is seen in most cases, with the exception of samples EPS_ISL_417413, EPS_ISL_421506, and to some extent EPS_ISL_425235. Additionally multimodal distributions are observed in several cases, which clearly suggest the presence of multiple equilibrium states for the N-finger. The most prominent peak is seen in sample EPS_ISL_423772, which corresponds to a larger amount of time spent away from a stable conformational equilibrium, even though a stable equilibrium was also visited (the lowest mode). While the mutation occurs on a beta strand located in a core area of the protein, the non-bonded interactions are similar for both M17 and I17. Out of 50 samples, only 10 sampled N-finger RMSD values similar to those of the reference chains. The rest displayed values of varying higher magnitude and duration displayed distributions suggestive of decreased stability. From this observation we can conclude that while the virus is still trying to evolve to past the human immune system, it could be accumulating mutations that can potentially make it less enzymatically active. However, with a relatively stable foundation in the 10 aforementioned samples (bearing mutations A255V, P99L, G15S, I136V, L232F, Y101C, A234V, V20L and T135I), it is possible that these may assist in perpetuating immune escape, without decreasing the enzymatic turnover rate.

**Fig. 4.**
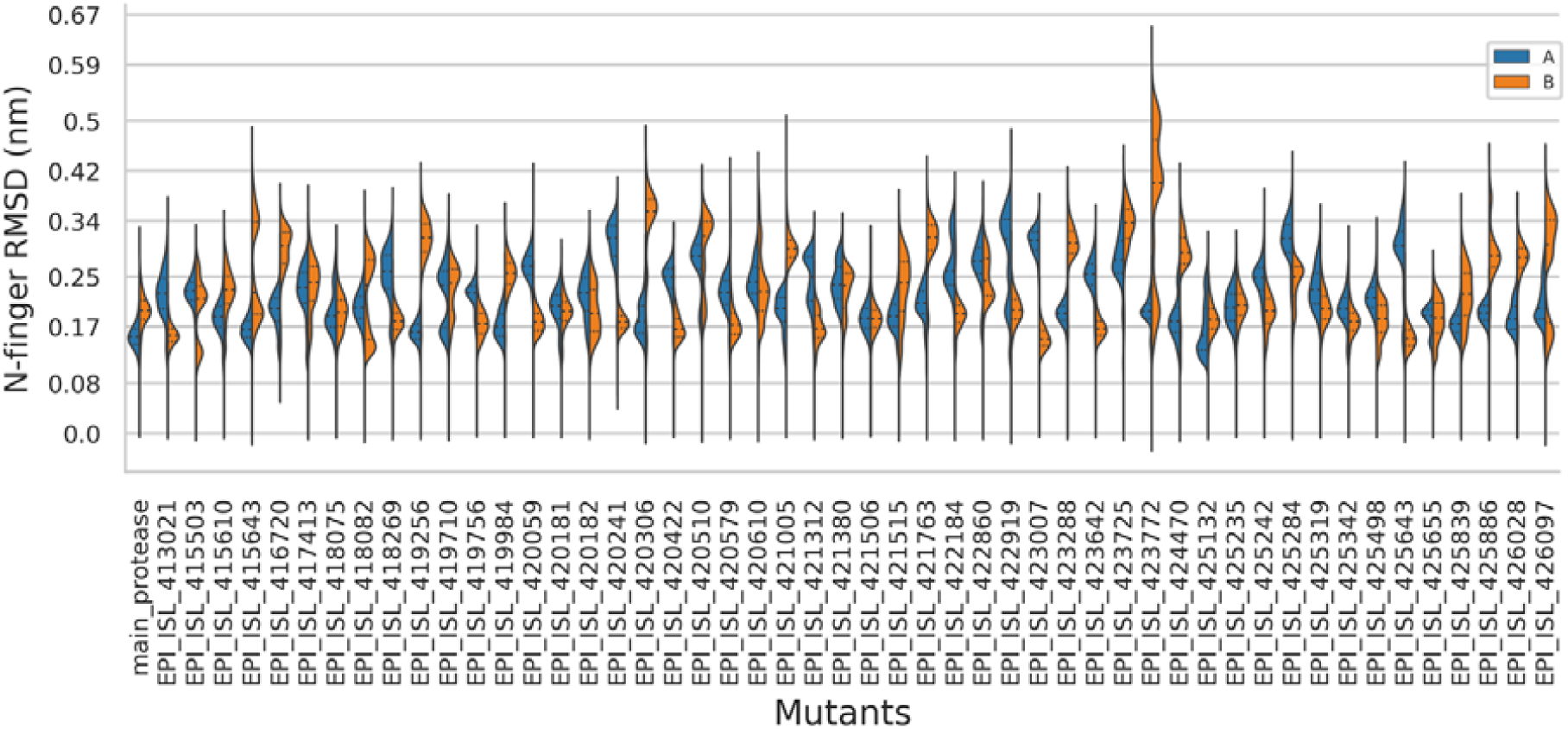
Kernel density distributions of RMSD values for the N-finger region across the mutant and reference protease complexes. The violin plots are split in two for each protein sample, showing the RMSD values for chains A (in blue) and B (in red). The tips of the distributions mark the minimum and maximum values for both chains combined in each complex.

### 2.5. Analysis of residue fluctuations and BC across M^pro^ samples

The analysis of residue fluctuations shows that there are generally interspersed areas of low and high flexibility along M^pro^ (Supplementary Fig. S1). Some local areas of higher flexibility were also seen, as is generally observed in unstructured secondary structures [54]. Additionally, from the lack of a clustering between chains belonging to the same sample for each segment of M^pro^ (Supplementary Fig. A1; cluster tree was removed for figure visibility), we conclude that the residue dynamics for the same domain between alternate chains of M^pro^ are asymmetric in the dimeric apo form. Focusing onto the specific regions of M^pro^, it can be seen that the N-terminal region of the N-finger generally displayed moderate residue fluctuations despite being sandwiched at the interface of the two M^pro^ chains, suggesting that it is not entirely immovable. Domain I is generally moderately flexible at residue positions 22-25, 33-34 and 92-93. Higher flexibility was globally observed along the intervals 45-65 and 71-74, and at position 76. Very high fluctuations were recorded for a subset of mutants at positions 46-54 most particularly for only one chain amongst the mutants EPI_ISL_416720, EPI_ISL_423642, EPI_ISL_415503 and EPI_ISL_425886, thus reinforcing the observation of asymmetry between chains. Mutations from samples EPI_ISL_416720 and EPI_ISL_425886 were already found to lead to increased backbone fluctuation using RMSD. The Y237H mutation in EPI_ISL_416720 introduces two carbon H-bonds (one with L272 and another with V237), in addition to the pi-alkyl interaction that is present in both the reference and this mutant, which seem to hold the solvent helices together in domain III. In a review by Horowitz and Trievel, the carbon H-bond was highlighted as an underappreciated interaction that is otherwise widespread in proteins, with the ability to form interactions as strong as conventional H-bonds via polarisation [55]. It is possible that such an increase in interaction may increase the stability around this region in domain III for the Y237H mutation. In the case of D248E mutation in EPI_ISL_425886, the D248 side chain is H-bonded to Q244 in the reference structure, possibly stabilising the helical structure. This interaction is absent upon mutating to an E248. In the last frame from MD, the side chain epsilon oxygen was found to interact with its backbone hydrogen atom, indicating a decreased stabilisation of the helical region of domain III. T201A in EPI_ISL_423642 abolishes the H-bond that is otherwise present between T201 and the backbone oxygen atom of E240, very likely weakening their interaction. The V157I mutation in EPI_ISL_415503 does not significantly alter the non-bonded interactions, but occurs on a beta strand on domain II. A unique behaviour was observed in the case of chain B of EPI_ISL_423772, where the leading residues of the N-finger were the most flexible while the rest of the protein was the least flexible across all samples. However, the M17I mutation does not significantly change the non-bonded interaction in EPI_ISL_423772, but occurs on a beta strand in domain I. Domain II is most flexible across all samples at residue position 153-155. Moderate flexibility is generally observed at residue positions 100, 107, 119, 137, 141-142, 165-171, 178 and 180. The linker region was generally highly flexible in all cases within the region spanned by residues 188-197. Notably higher fluctuations were observed at position 185 in samples EPI_ISL_420610 (chain B) and EPI_ISL_423007 (chain A). Across all domains, parts of domain III contain the most flexible residues within M^pro^. It is highly flexible at residue positions 222-224, 231-233, 235-236, 244-245 and 273-279. The high fluctuations observed C-terminus residues may not be very informative in our case as they freely interact with the solvent and do not have a strong enough network of non-bonded interactions with the M^pro^ domains. As a whole, from the empirical cumulative distribution (ECD) of averaged RMSF values across all M^pro^ samples, the top 5% positions (most variable regions) comprise residues 222, 277, 223, 154, 47, 72, 50, 224, 232, 64, 279, 236, 235, 244 and 51 ranging from 0.386 to 0.245 nm. The bottom 5% (most stable regions) of the distribution comprise positions 149, 146, 147, 174, 29, 113, 163, 39, 124, 150, 7, 175, 38, 161 and 173, ranging from 0.057 to 0.077 nm. On the 3D mapping (Fig. 5A), it can be seen that the regions with the highest flexibility are solvent exposed surfaces, comprising loops or parts of helices connected by loops. The central core of the enzyme has the lowest flexibility, most likely to provide structural stability to the functional dimer. Catalytic residues (HIS41 and CYS145) are connected to these stable core residues on one side. However, HIS41 is connected on the other side to a more mobile structure composed of a 3_10_ helix connected by loops on each end, which forms a lid structure, similar to what is described earlier for previous human coronavirus strains [56,57] 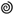.

**Fig. 5.**
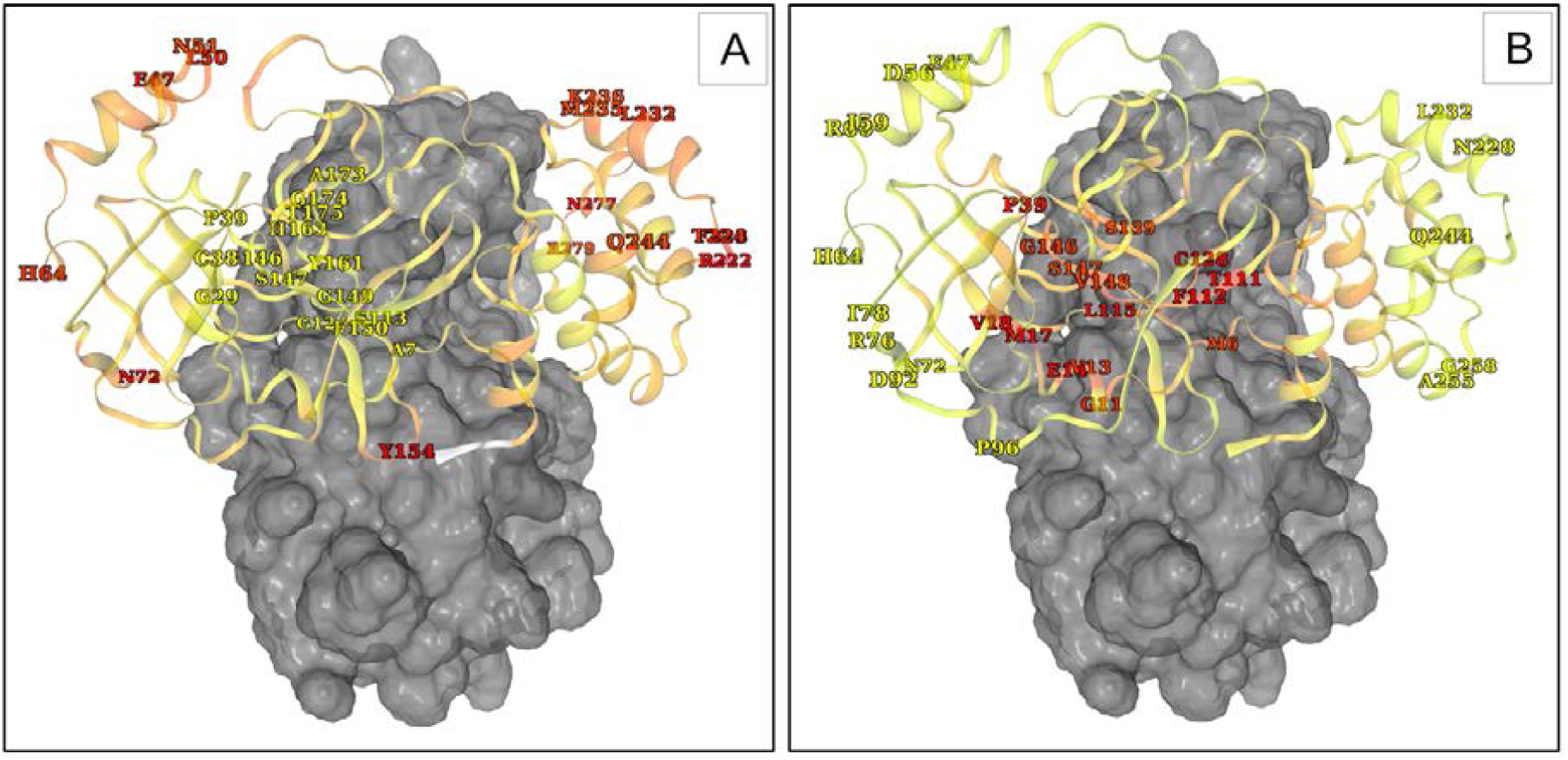
3D mapping of averaged values for (A) RMSF and (B) the average *BC*, computed across all Mpro samples. Only the extremes (top and bottom 5% of the ECD values across all samples) of averaged values are labelled for each metric. The lowest averaged values are in yellow while the highest ones are in red. The last three C-terminus residues (in white) were not mapped by RMSF as their high values would mask other values. While only one protomer is detailed, the data is applicable to both protomers. The other protomer is represented as a grey surface.

*BC* is a network centrality metric that is maximised when a large number of nodes (C_α_ or GLY C_β_ atoms) traverse a single node along geodesic paths to reach nodes within a network. When averaged from MD frames, this metric has been shown to be approximately inversely related to RMSF [58]. These values were very similar across all samples (Supplementary Fig. A2). For this reason, the discussion of our findings will be around the overall *BC* behaviour recorded across all samples. High and low *BC* values are present within all domains (Supplementary Fig. A2). The N-finger was found to be generally composed of high *BC* residues, most likely due to its high degree of non-bonded interactions between the protease chains. A large portion of domains I and III have low *BC* values, most likely due to the comparatively lower amount of contact with protein surfaces, which allows for a relatively higher mobility compared to domain II. The top 5% highest averaged *BC* values across all M^pro^ samples were either found in the monomer core regions or at the dimer interface, as seen in Fig. 5B. These comprised residue positions 17, 128, 115, 111, 112, 14, 18, 39, 11, 13, 146, 148, 147, 139, 6 and 140 in descending order of overall averaged *BC* values, ranging from 10480.987 to 6064.650. The residue positions within the lowest 5% overall averaged *BC* values comprise the positions 96, 72, 92, 258, 64, 244, 47, 59, 228, 56, 76, 60, 255, 78 and 232, ranging from 8.095 to 46.251. A complete listing of the top *BC* residues for each protomer is given in Table S2. Compared to the high *BC* areas that correspond to very well maintained communication residues (which may well be actual functional sites [58] the low *BC* values represent residues that are least important for maintaining the flow of communication across the protein, due to the transient nature of their short to medium ranged path lengths. From the computed sample average BC values and as seen the supplementary Table S2, residues positions 17 and 128 were found to occur as the most common first two residues in all cases. In the previous work done in human heat shock protein, high *BC* residues found within cavities were found to correspond to allosteric hotspots, as these had been independently verified by the sequential application of external forces on protein residues using the perturbation response scanning (PRS) method. In our case, however these two positions are not found in cavities, and are rather buried structural units that are not very mobile in the short to medium range. The high *BC* at M17 is possibly due to the increased stability imparted by the dimer interface. Due to the high centrality of these residues, it is possible that mutations leading to the alteration of their physicochemical activity may be accompanied by a decrease in the dimer stability.

### 2.6. Estimation of interprotomer distances using COM distance

The COM distance distributions between the M^pro^ protomers (Fig. 6) indicate that the distance between them can vary from over about 2.4 nm to 2.8 nm. Globally, which suggest the presence of at least two different equilibrium chain COM distances in these cases. More specifically, in isolates EPI_ISL_421763 and EPI_ISL_419984 a significant proportion of the sampled conformations depict the presence of closer chain COM values, though the percentage of such conformations is lower in EPI_ISL_419984. COM may be influenced to a certain extent by the shape of the domain arrangement and also plays a role in influencing the COM distance. Even though the distributions of interprotomer COM distances differ to varying degrees across samples, these differences may not be easily distinguished by visual inspection. Therefore, the next step was to quantify another aspect of the domain geometry, which is the inter-domain angle.

**Fig 6.**
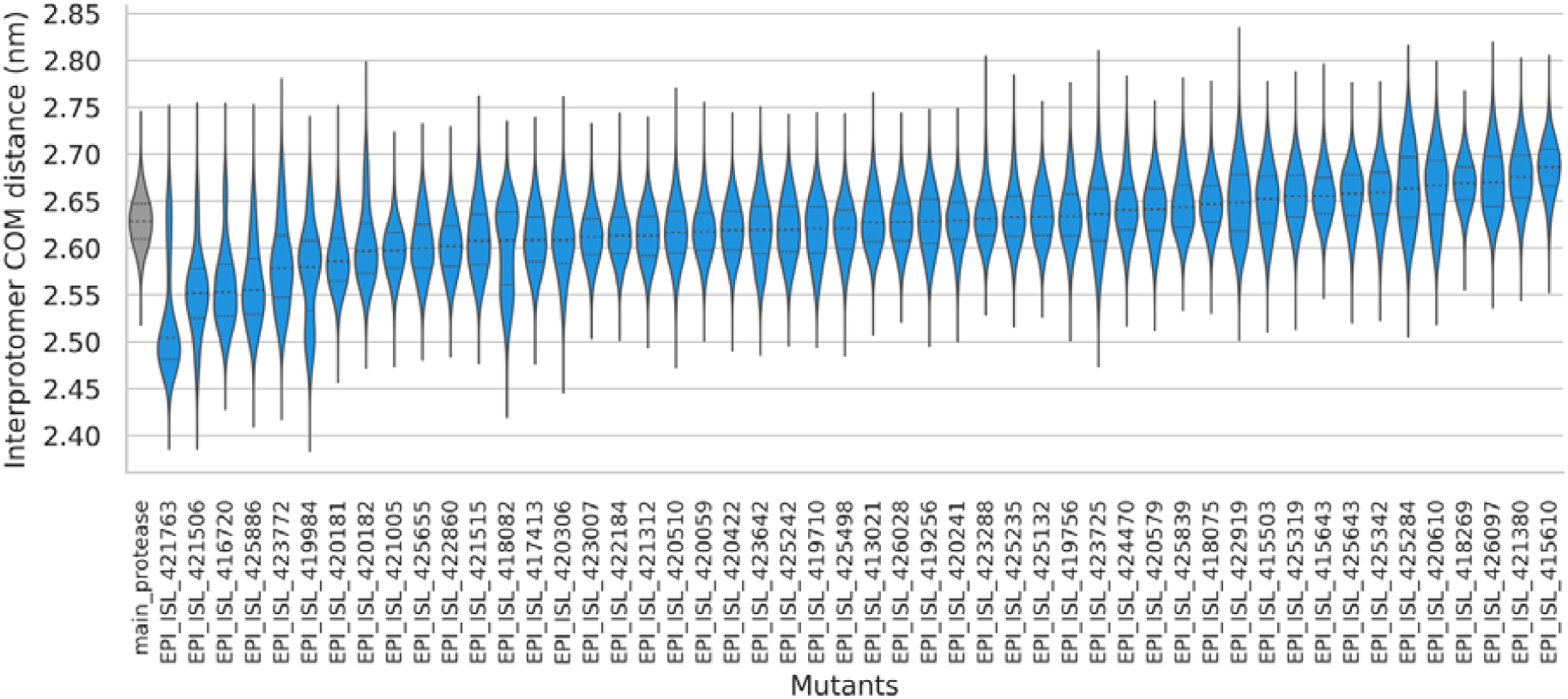
Distributions of interprotomer COM distances across samples, arranged in ascending order of average distance. The reference proteases are coloured in grey while the mutants are in blue.

### 2.7. Estimation of inter-domain angles in each protomer of M^pro^

The angle formed between domains I, II and II is has a relatively high information content for M^pro^ dynamics. While both chains A and B behave similarly in the reference, a higher degree of inter-domain angle variation is seen across the mutants, comprising skewed distributions and angle asymmetries between protomers (Fig. 7). The angle distributions are similar to those of the reference in several cases, however there are also many cases where angle distributions differ, as seen in the samples EPI_ISL_425643, EPI_ISL_421312, EPI_ISL_420241, EPI_ISL_418075, EPI_ISL_421005, EPI_ISL_419756, EPI_ISL_425342, EPI_ISL_420181, EPI_ISL_419256, EPI_ISL_422860, EPI_ISL_423007, EPI_ISL_425839, EPI_ISL_425132, EPI_ISL_425284, EPI_ISL_421515, EPI_ISL_413021, EPI_ISL_417413, EPI_ISL_419710, EPI_ISL_418269, EPI_ISL_426028, EPI_ISL_423642, EPI_ISL_425886 and EPI_ISL_416720. Additionally, among these samples, chains A and B were found to sample more divergent inter-domain angles, as seen from the shifted quartiles in samples EPI_ISL_420241, EPI_ISL_421005, EPI_ISL_419756, EPI_ISL_419256, EPI_ISL_422860, EPI_ISL_423007, EPI_ISL_425839, EPI_ISL_425132, EPI_ISL_425284, EPI_ISL_421515, EPI_ISL_413021, EPI_ISL_417413, EPI_ISL_419710, EPI_ISL_423642, EPI_ISL_425886 and EPI_ISL_416720. This provides additional support behind our general observation of the protomers generally behaving in an independent manner. It is possible that this independence might be of functional importance. Due to the richness of information retrieved from the measurement inter-domain angles, we propose that this metric may help assist the *in silico* characterisation (or differentiation) of M^pro^ variants from future strains of SARS-CoV-2 should any particular virus-associated phenotype become available.

**Fig. 7.**
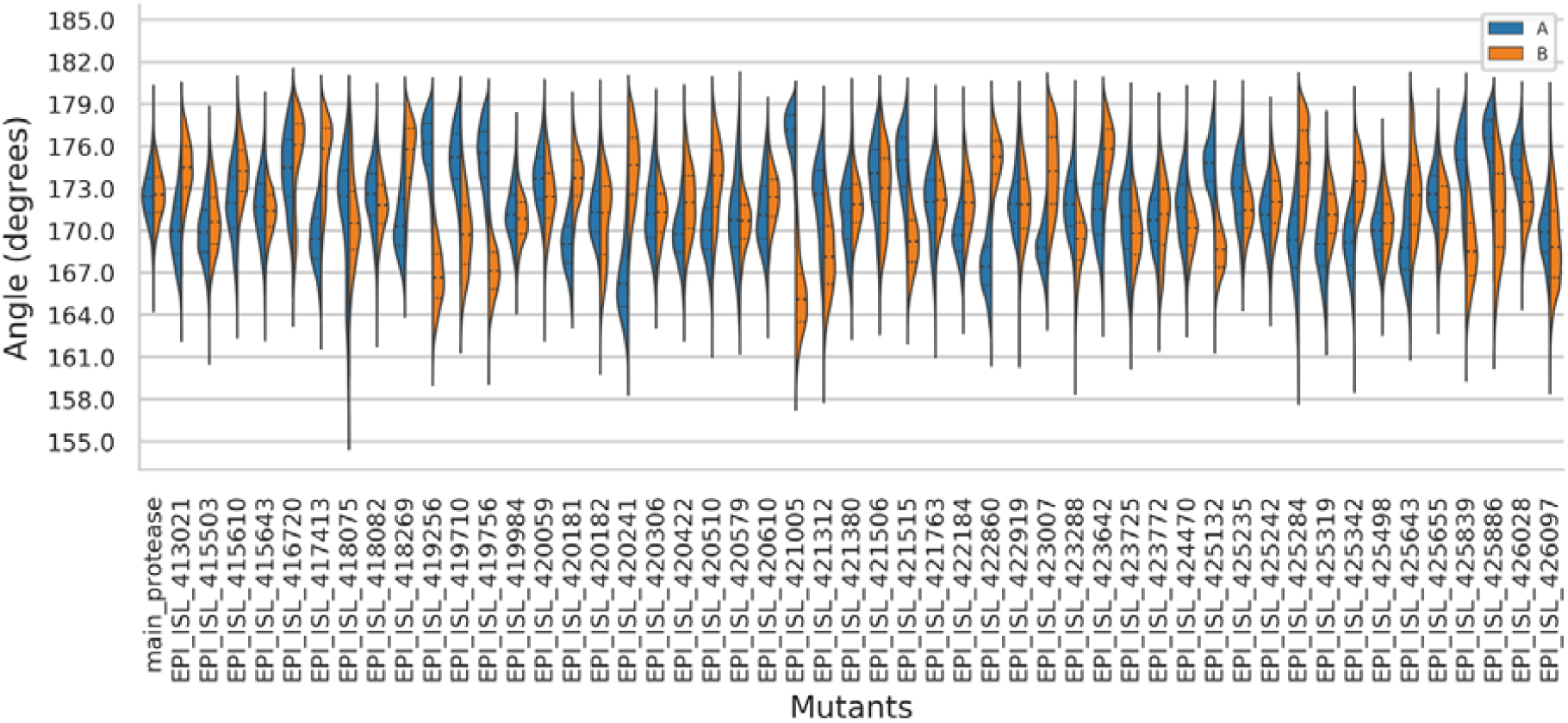
Kernel density distributions of inter-domain angles (domains I-II-III) across the mutant and reference protease complexes. The violin plots are split in two for each protein, showing the inter-domain angles for chains A (in blue) and B (in red). The tips of the distributions mark the minimum and maximum values for both chains combined in each protein complex.

### 2.8. Pocket detection and estimation of their compaction using R_g_

Pocket predictions from FTMap [59] and PyVOL [60] were concordant for all cases, except for the substrate binding site, which FTMap did not detect (Fig. 8). However, a very good coverage of interprotomer cavities was obtained by combining both methods, but more interestingly a potential allosteric site was found at the interprotomer interface, next to the substrate binding site. It was mirrored across each side of the interacting protomers. For this reason, the dynamics of both pockets were examined, as this could play an important role in the dimerisation properties of the protomers. The substrate binding site from each protomer was also examined due to its already known functional importance in catalysis. While the substrate binding pockets are easily defined as belonging to a given protomer, the interfacial pocket is composed of residues from each chain, namely residues 116, 118, 123, 124, 139 and 141 on chain A, and residues 5-8, 111, 127, 291, 295, 298, 299, 302, 303 on chain B. The mirrored interfacial pocket comprised the same residue positions, but for the opposite chain labels. The substrate binding site comprised residues 25, 26, 27, 41, 44, 49, 140, 141, 142, 143, 144, 145, 163, 166 in each protomer. The FTMap probes formed identical cross-clusters composed of probe IDs 1amn, 1ady, 1eth and 1acn at each site, which respectively correspond to the compounds methylamine, acetaldehyde, ethane and acetonitrile. As noted from the probe chemical compositions, this dual site has the potential to accommodate compounds of low molecular weight, with no (or probably limited) rotatable bonds and a low octanol-water partition coefficients [61].

**Fig. 8.**
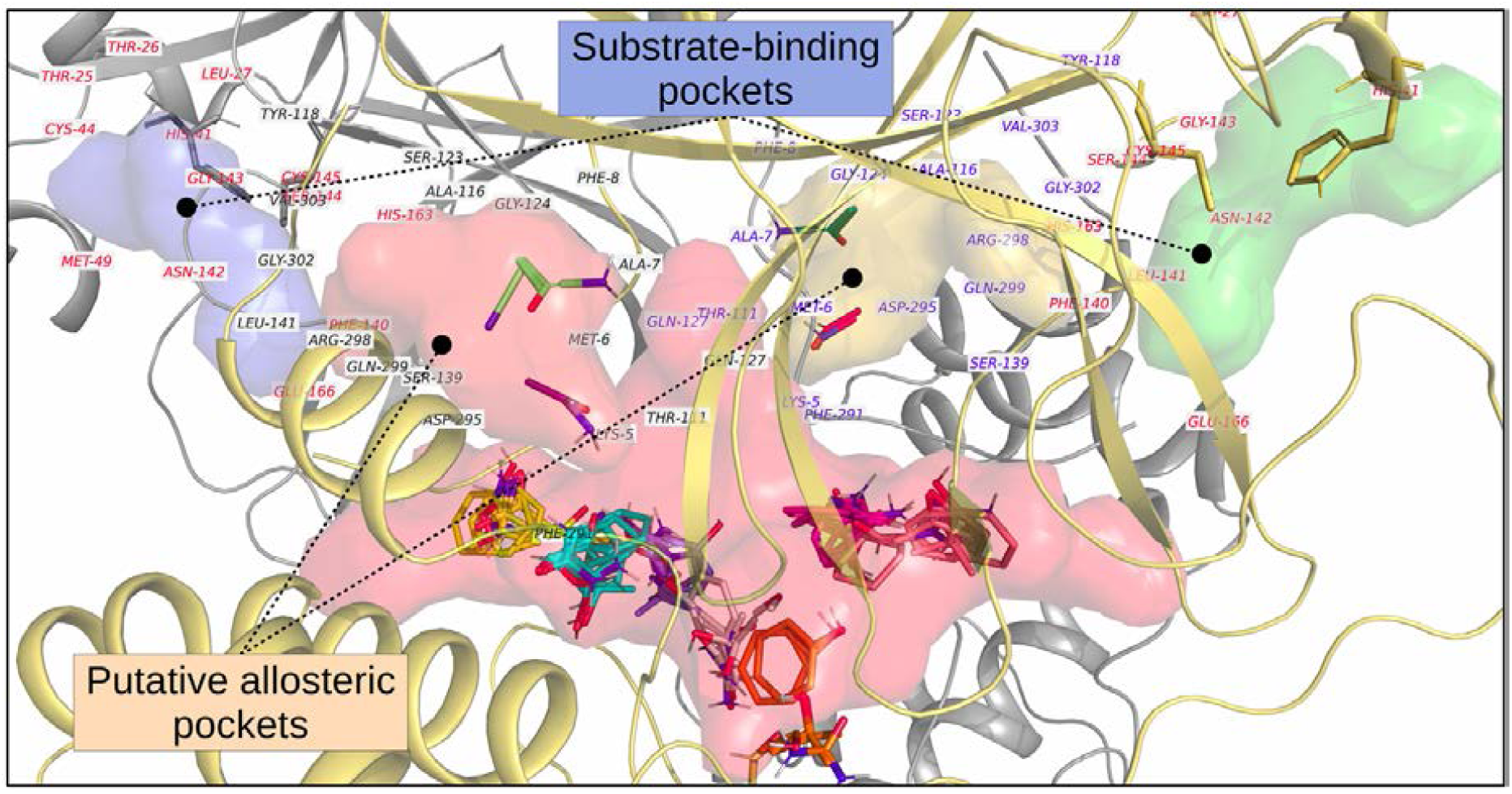
Pocket detection using combined predictions from FTMap and PyVOL. FTMap probes are shown as stick figure representations, while those from PyVOL are shown as surfaces. The protomers are depicted as cartoon representations, in grey and light orange.

From the distributions of R_g_ values for the catalytic site (Fig. 9.), we promptly observe the asymmetry between substrate binding pockets coming from each protomer in each sample. Partial symmetry is seen in few cases, such as EPI_ISL_419710, EPI_ISL_425242, EPI_ISL_423007, 416720, EPI_ISL_421380, EPI_ISL_425284 and EPI_ISL_419256. In the reference sample we observed a shifted in equilibrium R_g_, where one protomer oscillates around value while its counterpart also explores another. Multi-modal distributions indicate the presence of more than one equilibrium, which hints at cavity expansion and compaction movements. The most compact substrate binding cavities are observed in one of the protomers from sample EPI_ISL_425489, and to some extent samples EPI_ISL_418082 and EPI_ISL_423642.

**Fig. 9.**
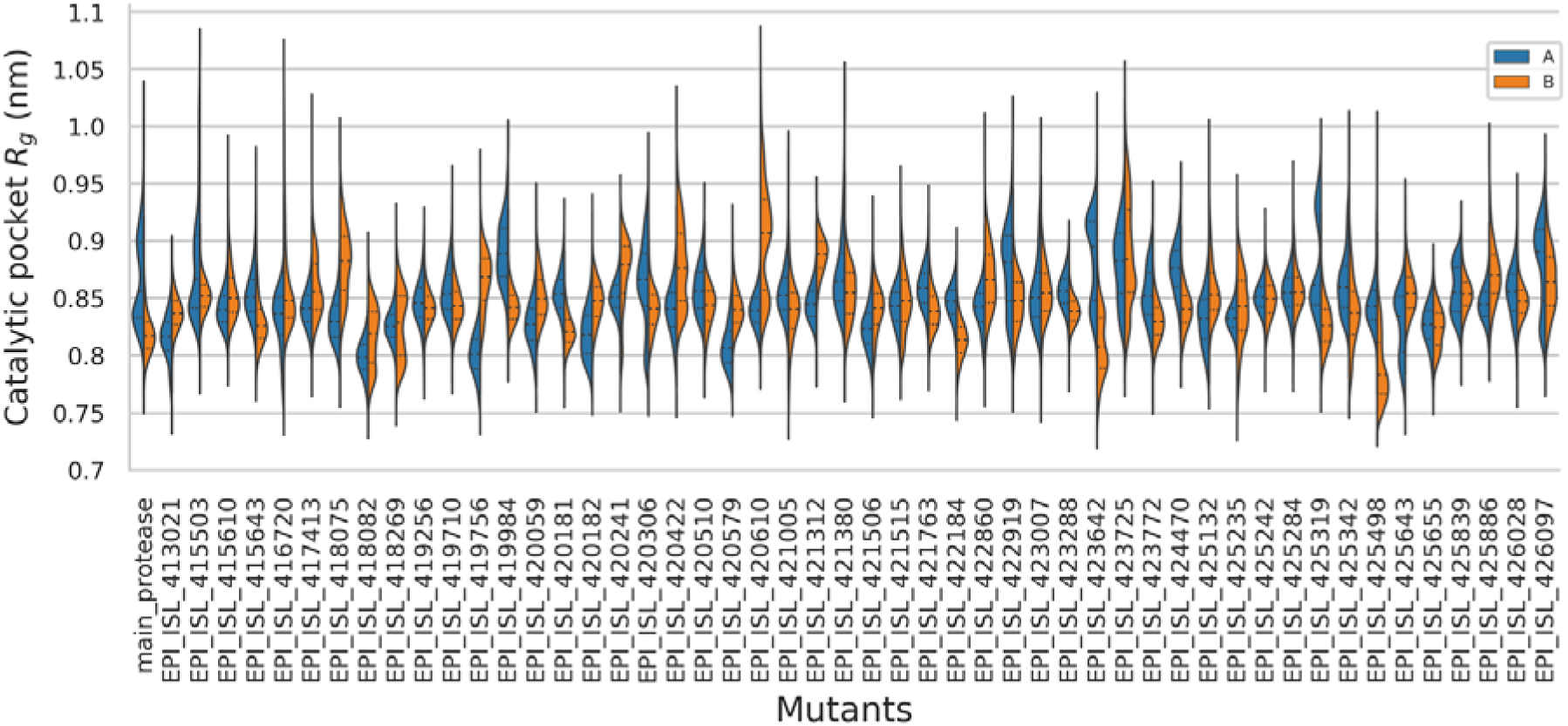
Kernel density distributions of R_g_ values for the substrate binding site from each protomer of Mpro. Chain A values are in red while chain B values are in blue. The maxima and minima are across protomers. Quartiles for each binding site are shown as dotted lines.

As done for the substrate binding cavity, the degree of compaction of the interprotomer cavities was also measured. Here as well, the distributions are asymmetric (Fig. 10). It is tempting to do a comparison between the substrate binding cavity and the interprotomer pockets. However, when the median R_g_ values obtained for the substrate binding pocket are correlated (using Spearman’s correlation) against the corresponding medians for the interprotomer pocket across all samples, no significant correlation was obtained, even though in our case, residue 141 was shared between both pockets. On an individual level, however there is a significant degree of correlation between the interprotomer pocket and the substrate binding site, ranging from -0.68 to 0.54 (using Spearman’s correlation), with p-values < 0.01 and absolute correlations > 0.1 for 64.9% of the sample comparisons (2 substrate binding site vs 2 interprotomer pockets for each sample). As the interprotomer pocket is mirrored between chains this may explain the negative correlations. From this finding we propose that the interprotomer pockets may play an important role in affecting the degree of compaction of the binding cavity and vice-versa. This suggests the possibility of a potentially bivalent modulation of the interprotomer pocket and the substrate binding site. As R_g_ only captures the overall degree of compaction, it may not entirely inform us about the cavity volume accessible to an allosteric modulator, however this may be an interesting lead for allosteric modulator targeting.

**Fig. 10.**
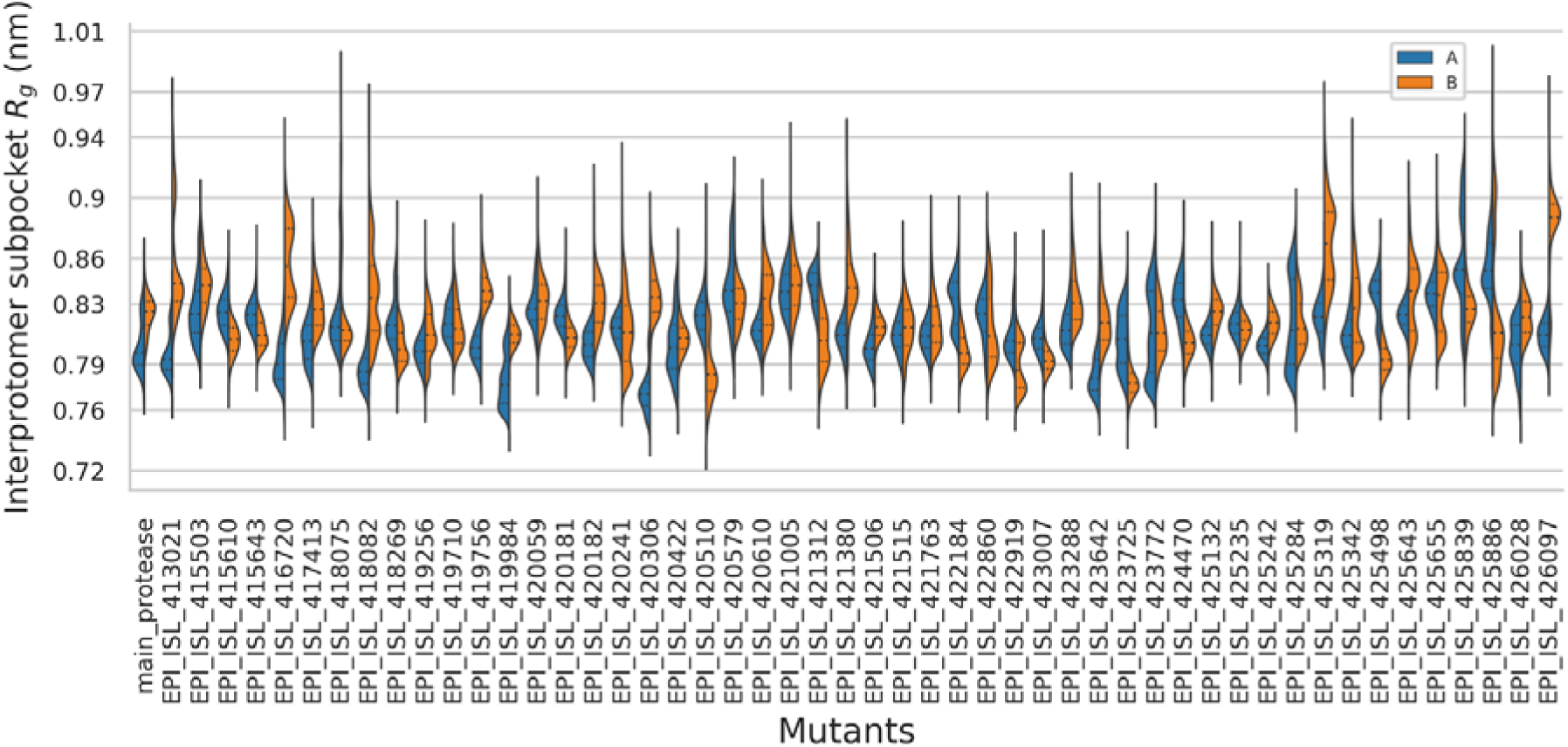
Kernel density distributions of R_g_ values across samples for the mirrored interfacial (and potentially allosteric) pockets.

### 2.9. Investigating correlated residue motions using Dynamic Cross Correlation

In order to compare the DCC across all samples, the DCC matrices were linearised before calculating pairwise correlations and clustering the final matrix (Fig. 11). While abstracting out the intricate details of residue correlation, clusters of correlated samples inform us on the global similarity of correlated protein motion across each pair of samples. From Fig 9. it can be seen that the samples are generally highly correlated, with the exception of samples EPI_ISL_415503, EPI_ISL_416720, EPI_ISL_419984, EPI_ISL_420241, EPI_ISL_422184, and EPI_ISL_418075, which form a sub-cluster of moderately correlated samples. Both EPI_ISL_415503 and EPI_ISL_416720 have been described in section 2.6 as having higher RMSF values at positions positions 46-54 due to the mutations V157I and Y237H. Mutation R105H (occurring on a loop region) in EPI_ISL_419984 leads to the loss of an H-bond with F181, but forms a pi-pi stacking interaction with Y182. The main difference in this case is in the higher protomer COM distance, as described in section 2.7. In sample EPI_ISL_420241, the P184L mutation occurrs in a solvent-exposed loop and no major non-bonded interactions were detected from the side chains or backbone atoms. The main reason for the observed difference is due to the increased divergence in inter-domain angles sampled from MD. In the case of EPI_ISL_422184, as explained in section 2.3, it was found the N-finger from one chains moved by a larger extent at the end of the simulation (∼93ns), diminishing its contacts with the alternate protomer, to interact more with its own protomer. This may be attributable to the S301L mutation, which reduces H-bonding at the end of the C-terminal helical structure. In the reference, four H-bonds are formed between S301, and the backbone oxygen atoms of V297 and R298, whereas two H-bonds are formed by L301. Sample EPI_ISL_418075 had the lowest correlation compared to all samples. Upon closer examination, it was found that the terminal alpha helix for each protomer was getting gradually destabilised towards the end of the simulation. This is very likely an artefact linked to the absence of the two C-terminal residues, as the residue is solvent exposed protein and display very similar residue interactions at the position 255 in both the reference and the mutant, even though A255V occurs on a helix.

**Fig. 11.**
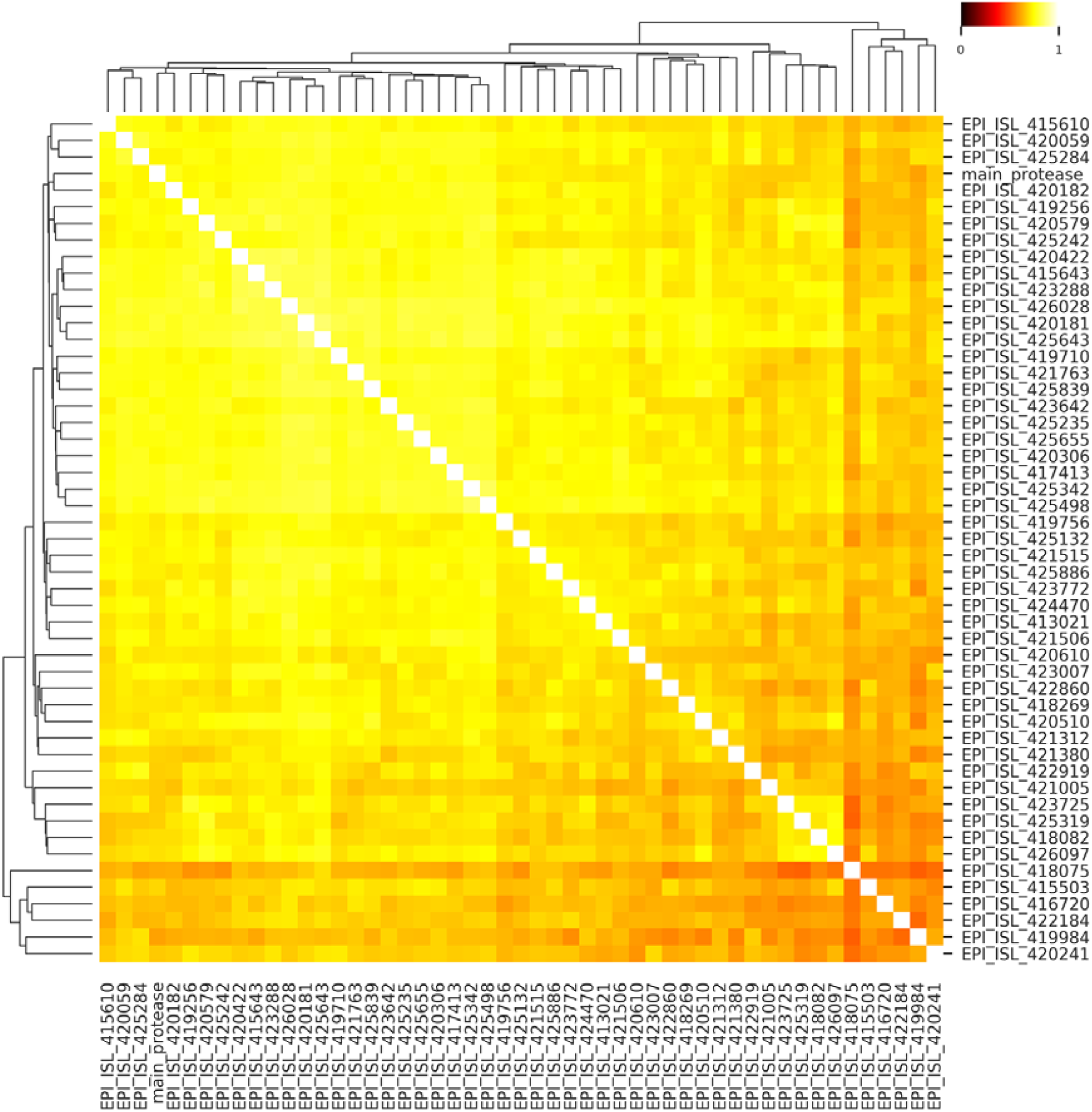
Heat map of correlations obtained from the linearised DCC matrices for the mutants and the reference M^pro^ clustered using the Euclidean distance.

A main observation across all samples is that residues within their individual domains are highly positively correlated within their respective domains. Domains I and II are seen to share a high degree of correlation with each other, behaving as a single unit most likely to maintain the integrity of the catalytic surface. From the simulated we find that domain III is generally not positively correlated with any of the other two domains, and can even be negatively correlated with itself on the alternate chain, and may suggest a degree of independence for domain III in terms of dynamics and possibly function as well, at least in its dimeric apo form.

### 2.10. Coarse-grained simulations of SARS-CoV-2 main protease structures reveal relationships between muta tional patterns and functional motions

To complement all-atom MD simulations and obtain a more granular description of structural dynamics in studied systems, we performed coarse-grained (CG) simulations of in the SARS-CoV-2 main protease structures in the free and ligand-bound forms using the CABS approach [46–50] (Fig. 12). By using a large number of independent CG simulations, we obtained conformational dynamics profiles for the studied systems and analysed these distributions in the context of examined mutations.

**Fig. 12.**
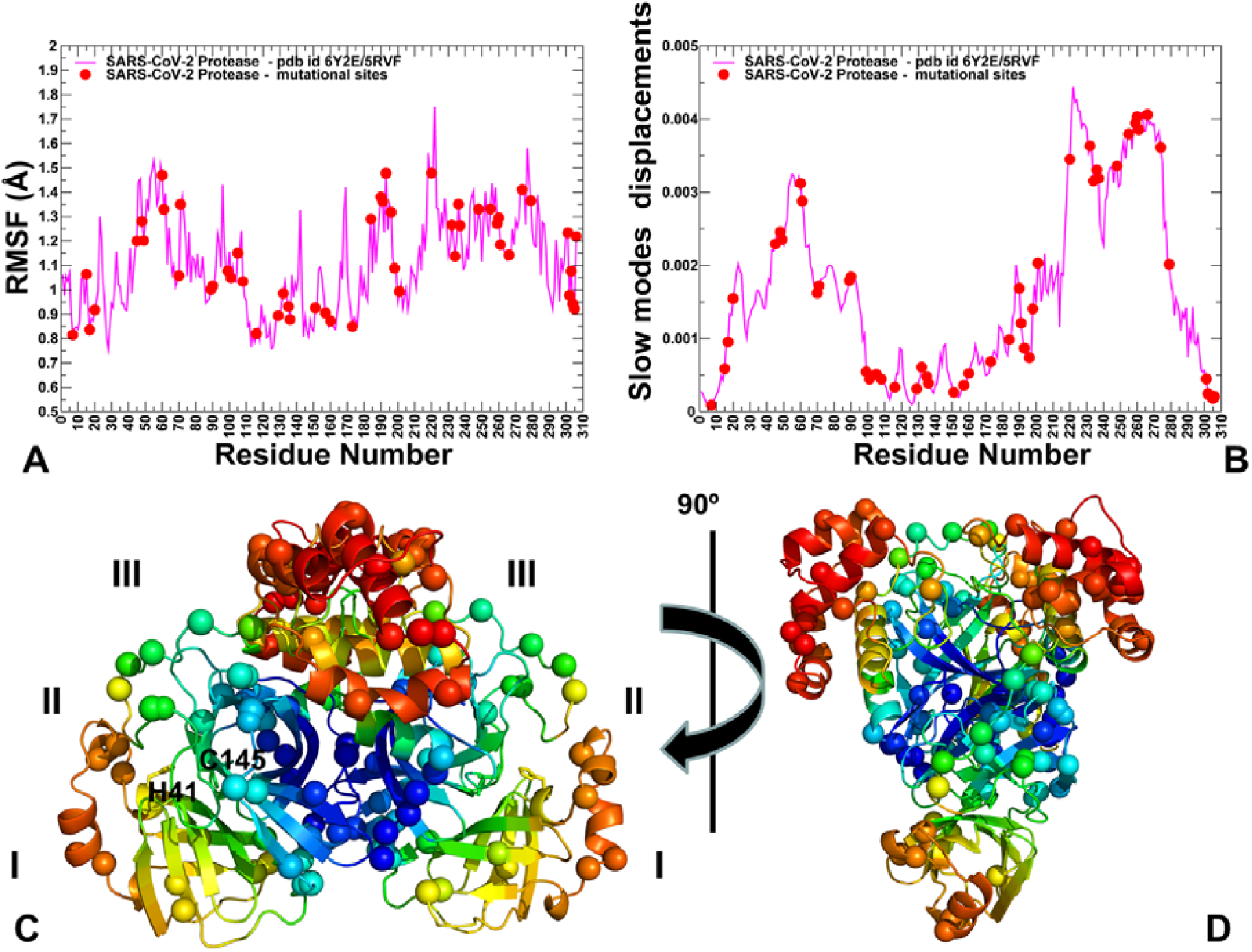
Conformational dynamics and collective motion slow mode profiles of the SARS-CoV-2 main protease structures (A) The computed root mean square deviations (RMSF) from CG MC dynamics simulations of the free enzyme of SARS-CoV-2 main protease (PDB ID: 5RVF, 6Y2E). The profile is shown in magenta lines. The positions of the residues undergoing mutations are shown in red filled circles (A7, G15, M17, V20, T45, D48, M49, R60, K61, A70, G71, L89, K90, P99, Y101, R105, P108, A116, A129, P132, T135, I136, N151, V157, C160, A173, P184, T190, A191, A193, T196, T198, T201, L220, L232, A234, K236, Y237, D248, A255, T259, A260, V261, A266, N274, R279 and S301L). (B) PCA analysis of functional dynamics SARS-CoV-2 main protease structures. The slow mode shapes are shown as mean square fluctuations averaged over first five lowest frequency modes (in magenta lines). The residues undergoing mutations are shown in red filled circles as in the panel A. (C) Structural map of the functional dynamics profiles derived from CG-MD simulations in the SARS-CoV-2 main protease (PDB ID: 5RVF, 6Y2E). The colour gradient from blue to red indicates the decreasing structural rigidity and increasing flexibility as averaged over the first five low frequency modes. The positions of the residues undergoing mutations are shown in spheres coloured according to their level of rigidity/flexibility in slow modes (blue-rigid, red-flexible). The locations of the protease domains I, II, and III are indicated. The catalytic residues HIS41 and CYS145 are shown in sticks. (D) The rotated view for the structural map of functional dynamics profiles in the SARS-CoV-2 main protease with sites of mutations in spheres.

RMSF of protein residues revealed the distribution of stable and flexible regions, thereby allowing the assessment of the extent of mobility for mutational sites (Fig. 12, panel A). In the domain I the most flexible residues are in the region 45-75, while for domain II the L2 loop (residues 165–172 and L3 (residues 185–200) located around the substrate-binding pocket also harbour a significant number of flexible positions. In addition, we observed significant fluctuations for surface residues 153–155 and 274–277 (Fig. 12, panel A). Of particular interest is the distribution of stable and flexible residues in the substrate binding pocket. Some pocket residues from domain I (T24, T25, T26, and L27) experience moderate fluctuations, while several other sites (M49, Y54) displayed considerably higher mobility. Another group of the substrate binding site residues from domain II (S139, F140, Q189) also exhibited appreciable fluctuations, while residues G143, S144, H164,H163, E166, P168 C145 showed only moderate changes and remained stable in CG simulations (Fig. 12, panel A). Notably and as expected the catalytic residues C145 and H41 from the substrate binding site also remained stable. The analysis generally showed that domain II residues were stable, while domain III (residues 198 to 303) showed more flexibility, especially in the peripheral solvent-exposed regions. This domain is involved in regulation of the dimerisation through a salt-bridge interaction between GLU290 of one protomer and ARG4 of the other protomer. Importantly, we found that these residues remained extremely stable in simulations. Interestingly, buried positions subjected to mutations exhibited different level of flexibility. While positions A7, V20, A116, A129, T135, I136, V157, C160, A173, and T201 were very stable, other buried sites with registered mutations in the domain III (A234, A266) showed larger fluctuations. It is worth mentioning that the interfacial residue A7 in the N-finger region important for enzymatic activity showed extreme level of rigidity.

Simulation-derived residue-residue couplings were evaluated using principal component analysis (PCA). By comparing slow mode profiles we found that functionally significant patterns can be yielded with up to the five slowest eigenvectors that account for ∼90% of the total variance of the dynamic fluctuations. The functional dynamics profile averaged over the five slow modes showed that the domain I (residues 10-99) and domain III (residues 198-303) are mostly mobile in functional motions and can undergo large structural changes (Fig. 12, panel B). At the same time, domain II (residues 100-182), is mostly stable during functional dynamics. The distribution of mutational sites clearly indicated the existence of two major clusters. One cluster of mutations is located in highly mobile regions of domain III (T198I, T201A, L220F, L232F, A234V, K236R, Y237H, D248E, A255V, T259I, A260V, V261A, A266V, N274D, R279C and S301L). These residues involved in protein motion are likely under different evolutionary constraints than are other functional sites. Another cluster of mutations is distributed in the domain II and includes 3 subgroups: a group of fully immobilized positions (A116V, A129V, P132L, T135I, I136V), a group of bridging (hinge-like) sites that connect rigid and flexible regions (Y101C, R105H, P108S, N151D, V157I/L, C160S, A173V, P184L/S) and a group of mostly mobile residues (T190I, A191V, A193V) (Fig. 12, panel B). The group of potential hinge sites may be important for controlling regulatory motions and mutations in these regions (such as V157I/L, P184L/S) may affect global movements in the protease and its enzymatic activity.

It is particularly important to dissect the connection between the function of some key residues and their contribution in collective movements. The dimerisation residues (R4, M6, S10, G11, E14, N28, S139, F140, S147, E166, E290, R298) are characterized by different local flexibility but tend to correspond to low moving regions of the protein in collective motions (Fig. 12, panel C, D). The key substrate binding residues (H163, H164, M165, E166, and L167) are located at the very border of structurally immobilized and more flexible regions, and as such may constitute a hinge region that controls cooperative movements. Notably, some other binding site residues D187, R188, Q189, T190, and A191 are more flexible in slow modes and may undergo functional motions. Substrate recognition sites tend to exhibit structural flexibility and sequence variations so as to enable specific recognition required for mediating substrate specificity. We also explored the functional dynamics profile of the ligand bound protease complex (Fig. 13). This structural map clearly illustrated that the ligand binding site is comprised of both rigid and flexible residues and located in the region that bridges area of high and low structural stability. In particular, we highlighted that residues S46, M49, T190, A191 in the substrate recognition site and in the ligand proximity may belong to moving regions in the global motions (Fig. 13, panel B).

**Fig. 13.**
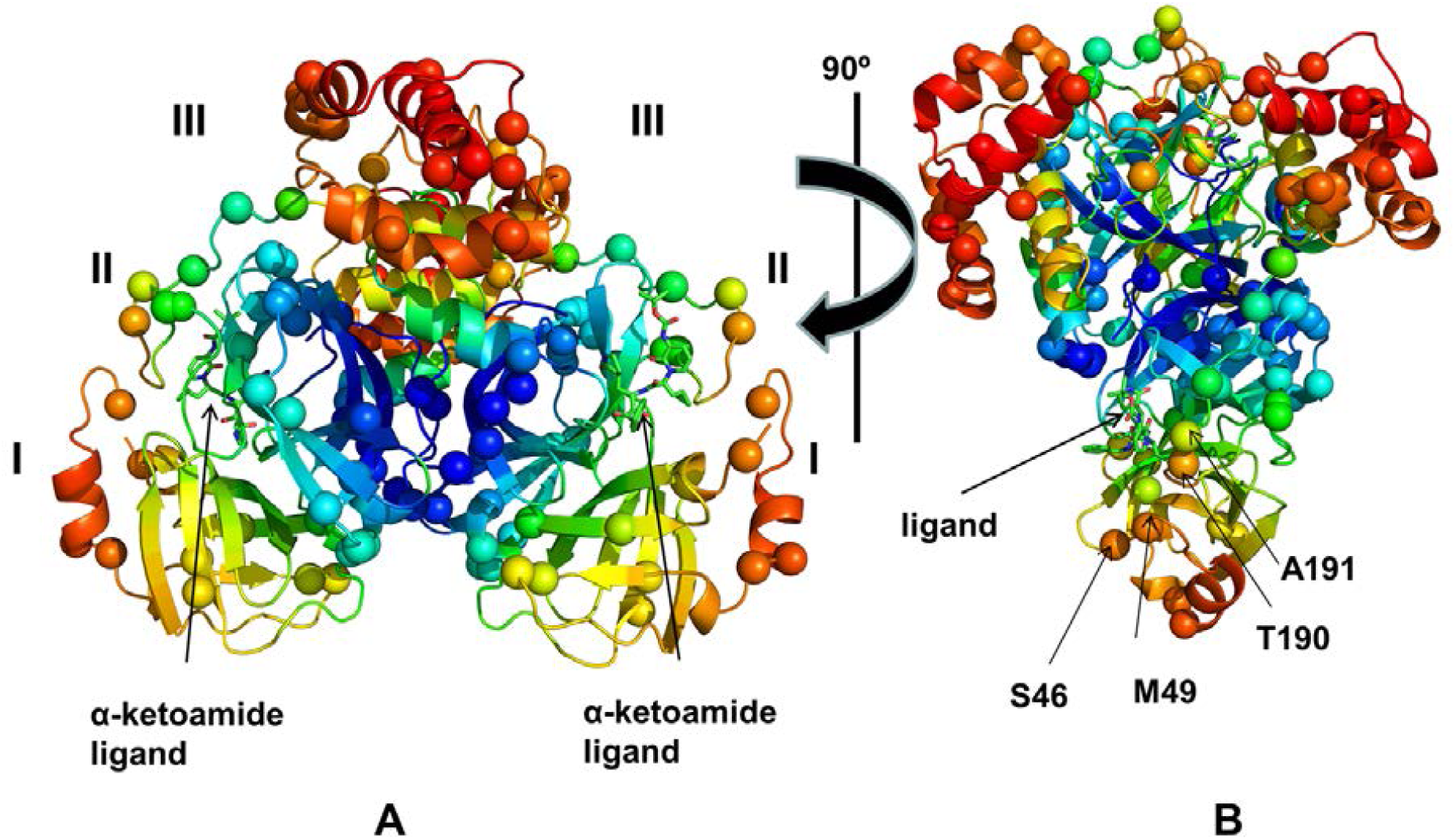
Structural map of functional motion profiles of the SARS-CoV-2 main protease structure complex with a ligand. (A) Structural map of the functional dynamics profiles derived from CG-MD simulations in the SARS-CoV-2 main protease in the complex with α-ketoamide ligand (PDB ID: 6Y2F). The slow mode shapes are averaged over first five lowest frequency modes. The colour gradient from blue to red indicates the decreasing structural rigidity and increasing flexibility as averaged over the first five low frequency modes. The positions of the residues undergoing mutations (A7, G15, M17, V20, T45, D48, M49, R60, K61, A70, G71, L89, K90, P99, Y101, R105, P108, A116, A129, P132, T135, I136, N151, V157, C160, A173, P184, T190, A191, A193, T196, T198, T201, L220, L232, A234, K236, Y237, D248, A255, T259, A260, V261, A266, N274, R279 and S301L) are shown in spheres coloured according to their level of rigidity and flexibility in the low frequency modes (blue-rigid, red-flexible). The locations of the protease domains I, II, and III are indicated. The α-ketoamide ligands are shown in sticks in both protomers. (D) The rotated view for the structural map of functional dynamics profiles in the SARS-CoV-2 main protease with sites of mutations in spheres. The position of the α-ketoamide ligand is shown in sticks. The mobile residues in the slow modes from the substrate binding site that form interactions with the ligand (S46, M49, T190, A191) are indicated by arrows and annotations.

Our analysis shows that structural clusters of mutations may be distinguished by their evolution propensity and global mobility in slow modes regions. The mobile residues may be predisposed to serve as substrate recognition sites, whereas residues acting as global hinges during collective dynamics are often supported by conserved residues. The observed conservation and mutational patterns may thus be determined by functional catalytic requirements, structural stability and geometrical constraints, and functional dynamics patterns. We found that some sites and corresponding mutations may be associated with dynamic hinge function. The mutability of hinge sites (Y101C, R105H, P108S, V157I/L, C160S, A173V, P184L/S) and nearby sites (T190I, A191V, A193V) may be related with their structural and dynamic signatures to reside in the exposed protein regions rather than in the more conserved protein core. We could also conclude that these sites are located near the active site and control in the bending motions needed for catalysis, so their mutability may have an important functional role for enzymatic activity especially when combined with mutations in adjacent regions.

## 3. Materials and Methods

### 3.1. Sequence and template retrieval

A high resolution (1.48 Å) biological unit for crystal structure of the SARS-CoV-2 main protease (PDB ID: 5RFV [62]) was retrieved from the Protein Data Bank (PDB) [63] to be used as a template for homology modelling. Its sequence was used as reference throughout this work. PyMOL (version 2.4) [64] was used to remove any non-protein molecule and to reconstitute the biological unit as chains A and B. SARS-CoV-2 genomes of any length were acquired from the GISAID website as a FASTA-formatted file [9] 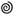.

### 3.2. Building the mutation data set

Low coverage sequences (entries with >5% unknown nucleotides) from GISAID were not selected. A local BLAST database was then set up for these sequences using the *makeblastdb* command available from the BLAST+ application (version 2.8.1) [65]. Protease mutants were subsequently retrieved using the reference sequence as query parameter for the *tblastn* command with default parameters, except for the maximum number of target sequences, which was set at 10000. Identical sequences were then filtered out before selecting BLAST hit sequences that had a 100% sequence coverage and a percentage sequence identity of < 100%. Sequences coming from non-human hosts were discarded. In order to retain as much data as possible and minimize the incorporation of sequencing errors, fold coverage was used where provided, while quality was imputed on the basis of an identical M^pro^ sequence being present more than once in the filtered dataset, unless a sequencing fold coverage value was available. This resulted in a 50 mutant sequences with either high coverage, where available or with additional sources of support otherwise. Further details about the sequences are given in Supplementary acknowledgement Table S1.

### 3.3. Homology modelling, pH adjustment and analysis of residue interactions

PIR-formatted target-template sequence alignment files were generated for each mutant using the BioPython library (Version 1.76) [66] within ad hoc Python scripts for use in MODELLER (version 9.22) [67]. The automodel class was used with slow refinement and a deviation of 2 Å to generate 12 models in parallel for each mutant, after which the ones with the lowest z-DOPE scores were retained. The protein was then adjusted to a pH of 7 using the PROPKA algorithm from the PDB2PQR tool (version 2.1.1) [68]. For visualising the overall interactions at given residue positions, the Arpeggio tool [69] was used to programmatically generate the inter-residue interactions, before computing their sums using an in-house Python script. More generally, the Discovery Studio Visualizer (version 19.1) was used for describing the non-bonded interactions [70].

### 3.4. Molecular dynamics simulations

All-atom protein MD simulations were run for the protonated dimers using GROMACS (version 2016.1) [71] at the Center for High Performance Computing (CHPC). Proteins were placed in a triclinic box containing 0.15M NaCl in SPC modelled water. A minimum image distance of 1.5 nm between the solute and the box was used. The system was then energy minimized using the steepest descent algorithm with an initial step size of 0.01 nm for a maximum force of 1000 kJ/mol/nm and a maximum of 50000 steps. Temperature was subsequently equilibrated at 310 K for 50 ps, according to the NVT ensemble. Pressure was then equilibrated at 1 bar for 50 ps, using the Berendsen algorithm according to the NPT ensemble. During both NVT and NPT, the protein was position restrained and constraints were applied on all bonds. 100 ns unrestrained production runs were then performed, with constraints were applied only on H-bonds and the Parrinello-Rahman algorithm was used for pressure coupling. In all cases a time step of 2 fs was used, with a short-range non-bonded cut-off distance of 1.1 nm and the PME algorithm for long-range electrostatic interaction calculations. Prior to analysis, the periodic boundary conditions were removed and the trajectories were corrected for rotational and translational motions.

### 3.5. Coarse-Grained Simulations

Coarse-grained (CG) models enable simulations of long timescales for protein systems and assemblies, and represent a computationally effective strategy for adequate sampling of the conformational space while maintaining physical rigour. The CABS model was employed for multiple CG simulations [46–50] of the SARS-CoV-2 main protease dimer structures (PDB ID: 5RFV, 6Y2E, and 6Y2F [29]). In this model, the CG representation of protein residues is reduced to four united atoms. The residues are represented by main-chain α-carbons (Cα), β-carbons (Cβ), the COM of side chains and another pseudoatom placed in the centre of the Cα-Cα pseudo-bond. The sampling scheme involved Monte Carlo (MC) dynamics moves including local moves of individual residues and moves of small fragments composed of 3 protein residues. 100 independent CG simulations were carried out for each studied system with the CABS-flex standalone Python package for fast simulations of protein dynamics, which is implemented as a Python 2.7 object-oriented package [50]. In each simulation, the total number of cycles was set to 1,000 and the number of cycles between trajectory frames was 100. Accordingly, the total number of generated models was 2,000,000 and the total number of saved models in the trajectory used for analysis was 20,000. It was previously shown that the CABS-flex approach can accurately recapitulate all-atom MD simulations on a long timescale [46–50]. The results of 100 independent CG-CABS simulations for each system were averaged to obtain adequate sampling and ensure convergence of simulation runs.

### 3.6. Dynamic Residue Network (DRN) analysis and Dynamic Cross Correlation (DCC)

The MD-TASK tool kit [34] was used to calculate the averaged *betweenness centrality (BC)* over the last 50 ns of simulation for each proteins using a cut-off distance of 6.70 Å and a step size of 25, generating a total of 10,001 frames. DCC was calculated for each of the proteins using the same frames and time step, before linearising each matrix. Pairwise Pearson correlations were then performed for all linearised matrices before performing hierarchical clustering. In all cases, the Cβ and glycine Cα atoms were used. The GROMACS commands *trjconv* and *make_ndx* were used to reduce the trajectory sizes, to only keep Cα and Cβ atoms prior to computation.

### 3.7. Pocket detection and dynamic analysis

The reconstituted biological unit for the reference structure was submitted to the FTMap web server using default parameters. The PyMOL plugin PyVOL was then used to identify the surfaces of any potential cavity, specifying the protein as selection, with the default minimum volume of 200 Å^3^. Predictions from both tools were combined. Residues for the interprotomer subpocket were defined by visually inspecting residues in close proximity to the cavity surfaces detected by PyVOL that overlapped with part of the FTMap probe binding predictions. The pocket was selected due to its location and accessibility to the outside. The substrate binding residues (identified using PyVOL) from both protomers of M^pro^ were also investigated due their functional importance in catalysis, even though FTMap did not identify a binding hotspot at that location. The radius of gyration (R_g_) for the subpocket and the binding pockets was then computed from the entirety of the simulated MD data in each case, using the GROMACS *gyrate* command. The generated data was then visualised and analysed using various open source Python libraries, such as matplotlib [72], Seaborn, Pandas [73], NumPy [74], SciPy [75], MDTraj [76] and NGLview [77].

## 4. Conclusions

COVID-19 represents a significant global threat for which no effective solution currently exists, with the exception of social distancing, which has slowed down the viral progression. Time is of the essence to find a cure that will counter the impact of the virus, especially for those at higher risk and for the world economies. Analysing the structural and dynamic properties of the novel mutants of the SARS CoV-2 M^pro^ gives important pointers and details about its dynamics behaviour.

From this work, several non-synonymous mutations were found across all domains of the SARS-CoV-2 M^pro^. There are various single residue substitutions, among which several are substitutions of alanine and valine. These mutations have occurred both in buried and solvent-accessible surfaces. From our filtered data set, residue positions 15, 157 and 184 appear to have mutated more than once. A relatively high number of titratable amino acids present in M^pro^, which we presume may play an important role in influencing its behaviour at various pH levels. Higher backbone flexibility was observed for the isolates EPI_ISL416720, EPI_ISL426097, EPI_ISL421763, EPI_ISL420610, EPI_ISL425284, EPI_ISL421380, EPI_ISL423772 and EPI_ISL425886. More importantly, a high number of samples displayed various levels of stability of the N-finger region, suggesting an active viral adaptation in the human host, trying to find a trade-off between viral fitness and immune egress. More generally, regions of lowest flexibility (and high *BC*) were core residues, while solvent-exposed loops were most flexible. All samples displayed a slight interprotomer twisting motion. A high degree of variation was observed in (1) the angle formed between domains I, II and II, (2) the substrate binding pocket R_g_, (3) the interprotomer pocket R_g_ and (4) the N-finger flexibility, which may all be good descriptors for characterising M^pro^ dynamics. *BC* values were very similar across all samples, with extreme values being essentially anti-correlated to RMSF. Residues 17 and 128 appear to be very central residues, and based on the *BC* network metric, it is likely that mutations altering their physicochemical property may have the potential to alter dimer stability. We propose the presence of a mirrored allosteric interprotomer pocket, supported by multiple cavity detection approaches, and correlations between the interprotomer pocket compaction and the substrate binding pocket. The mirrored pocket may have the potential to accommodate compounds of low molecular weight and polarity. Asymmetries to partial symmetries in R_g_ distributions were seen for each of the substrate binding pockets and the interprotomer cavities for each isolate. However, a large portion of the samples displayed overall positive correlations according to DCC. In each individual DCC plot, domains I and II are found to behave as a single unit while domain III is generally more independent. Thorough, independent CG simulations of the apo and the ligand-bound M^pro^ further revealed a connection between regions accumulating clusters of mutations and their degree of residue fluctuation from the slowest modes. Additionally, we report of a possible set of dynamic hinging residues and their tendency to acquire mutations in exposed protein regions, whilst being grounded by less mutable core residues.

As a final note, it is important to be aware that there is an inherent lack of sampling depth in this current analysis of COVID-19 sequences due to the existence of undiagnosed mutations that may be present among infected individuals [78] at the time of writing, and in our case we might have missed certain mutations using our filtering criteria, in our effort to balance accuracy and the number of high confidence representative samples. Therefore frequencies should be handled with caution especially at the early stages of the SARS-CoV-2 evolution.

## Supplementary Materials

**Fig. S1.** Heat map for the RMSF recorded from each chain of the Mpro samples. The data has been clusterd separately for each segment of the protase (shown along the y-axis). The dendrograms have been removed for clarity. RMSF are plotted separately to highlight domain level differences, which have their own scale. Domains I-III are annotated as red, blue and orange strips respectively. The N-finger is in cyan while the linker is coloured green. Active site residues are denoted by the letter “x”. **Fig. S2.** Heat map of averaged BC values for each chain of the Mpro samples. Each sample has been clustered using hierarchical clustering with the Eucldean distance metric. Domains I-III are annotated as red, blue and orange strips respectively. The N-finger is in cyan while the linker is coloured green. Active site residues are denoted by the letter “x”. **Table S1**. Acknowledgement and sequence support. **Table S2.** High *BC* residues are shown for each chain (A and B).

## Author Contributions

Conceptualization, Olivier Sheik Amamuddy and Özlem Tastan Bishop; Data curation, Olivier Sheik Amamuddy; Formal analysis, Olivier Sheik Amamuddy, Gennady M Verkhivker and Özlem Tastan Bishop; Methodology, Olivier Sheik Amamuddy and Gennady M Verkhivker; Resources, Özlem Tastan Bishop; Software, Olivier Sheik Amamuddy; Supervision, Özlem Tastan Bishop; Visualization, Olivier Sheik Amamuddy and Gennady M Verkhivker; Writing – original draft, Olivier Sheik Amamuddy; Writing – review & editing, Olivier Sheik Amamuddy, Gennady M Verkhivker and Özlem Tastan Bishop.

## Funding

This research received no external funding.

## Acknowledgements

We would like to thank the Centre for High Performance Computing for providing the computer time for the simulations. We also thank GISAID making available the COVID-19 genomic material, the originating and submitting laboratories (acknowledged in Supplementary Table A1). Genetic sequences are purposely not disclosed in this article due to the restrictions from GISAID, but are accessible from the database.

## Conflicts of Interest

The authors declare no conflict of interest.

## Abbreviations

*BC*: *Betweenness centrality*
BLAST: Basic Local Alignment Search Tool
CG: Coarse-grained
CHPC: Centre for High Performance Computing
COM: Centre of mass
DCC: Dynamic Cross Correlation
GISAID: Global Initiative on Sharing All Influenza Data
PDB: Protein Data Bank
PME: Particle Mesh Ewald
MC: Monte Carlo
MD: Molecular dynamics
RMSD: Root mean squared deviation
RMSF: Root mean squared fluctuation
COVID-19: Coronavirus disease 2019
SARS-CoV: Severe acute respiratory syndrome coronavirus
SARS-CoV-2: Severe acute respiratory syndrome coronavirus 2

## Notes

### Competing Interest Statement

The authors have declared no competing interest.

