## Supplementary figure for "Impact of emerging mutations on the dynamic properties the SARS-CoV-2 main protease: an *in silico* investigation"

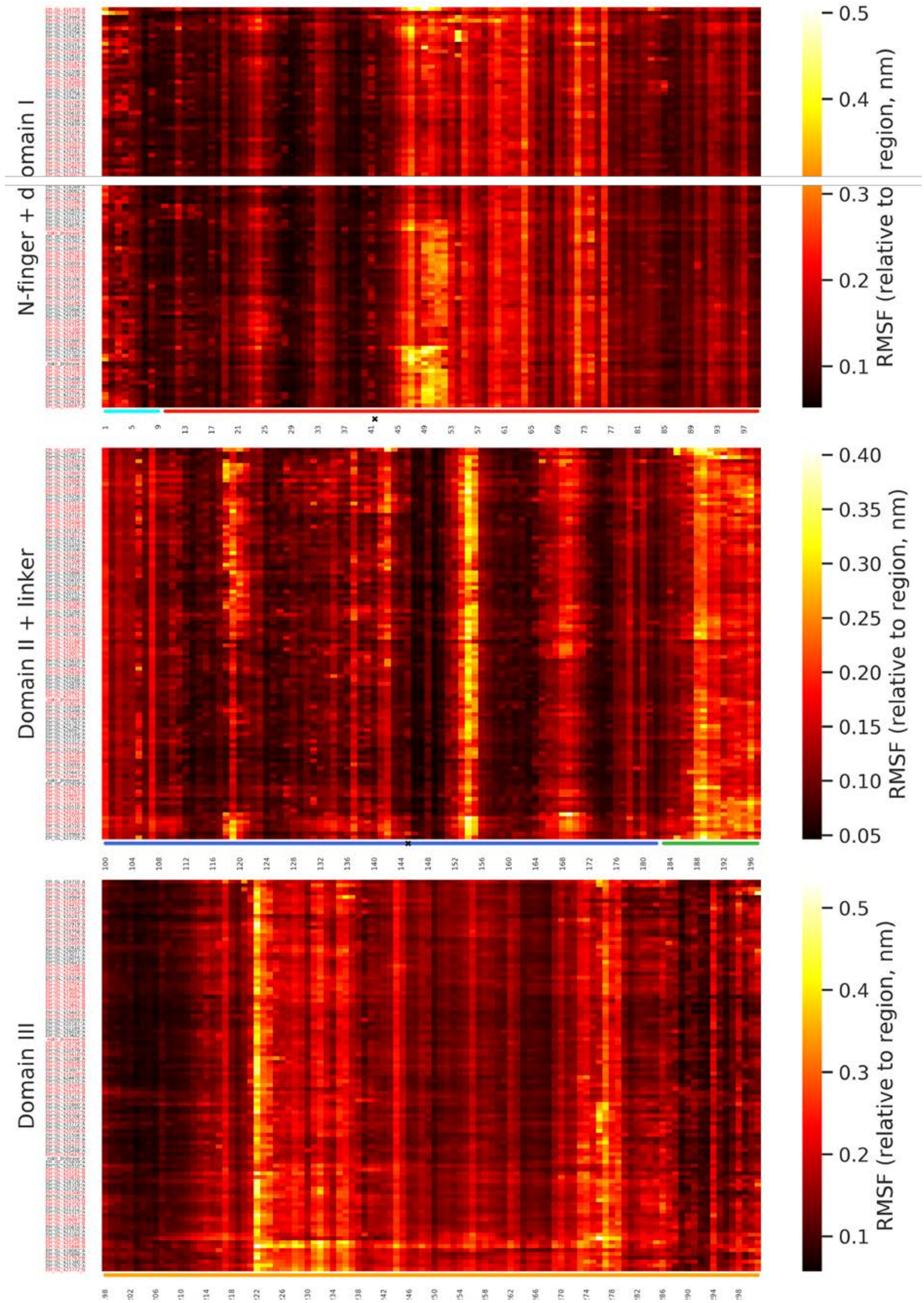


Fig. S1. Heat map for the RMSF recorded from each chain of the M^pro^ samples. The data has been clusterd separately for each segment of the protase (shown along the y-axis). The dendrograms have been removed for clarity. RMSF are plotted separately to highlight domain level differences, which have their own scale. Domains I-III are annotated as red, blue and orange strips respectively. The N-finger is in cyan while the linker is coloured green. Active site residues are denoted by the letter "x".

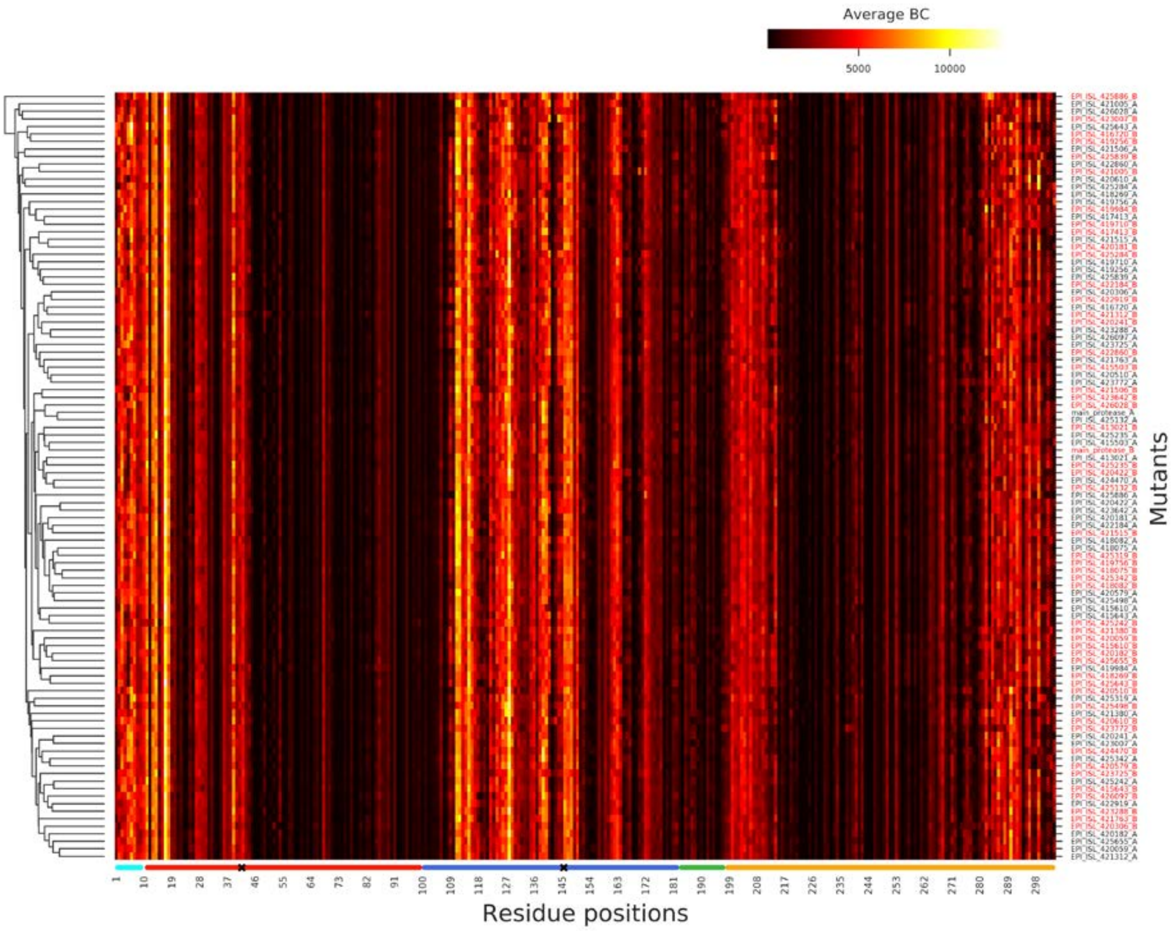


Fig. S2. Heat map of averaged BC values for each chain of the M$^{pro}$ samples. Each sample has been clustered using hierarchical clustering with the Eucldean distance metric. Domains I-III are annotated as red, blue and orange strips respectively. The N-finger is in cyan while the linker is coloured green. Active site residues are denoted by the letter "x".
