## Supplementary table for "Impact of emerging mutations on the dynamic properties the SARS-CoV-2 main protease: an *in silico* investigation"

### Title

**Supplementary Table A1: Acknowledgement and sequence support**

| Accession number | Virus name | Sequence support | Originating Lab | Submitting Lab | Authors |
|---|---|---|---|---|---|
| EPI_ISL_413021 | hCoV-19/Switzerland/1000477757/2020 | 444x | Klinik Hirslanden Zurich | Institute of Medical Virology, University of Zurich | Stefan Schmutz, Maryam Zaheri, Verena Kufner, Gabriela Ziltener, Patrick Redli, Fiona Steiner, Jon Huder, Riccarda Capaul, Andrea Zbinden, Jürg Böni, Michael Huber, Christian Ruef, Alexandra Trkola |
| EPI_ISL_415503 | hCoV-19/Netherlands/NoordBrabant_46/2020 | 30x | Dutch COVID-19 response team | Erasmus Medical Center | David Nieuwenhuijse, Bas Oude Munnink, Reina Sikkema, Claudia Schapendonk, Irina Chestakova, Anne van der Linden, Mark Pronk, Pascal Lexmond, Corien Swaan, Manon Haverkate, Madelief Mollers, Mart Stein, Sandra Kengne Kamga Mobou, Jeroen van Kampen, Jolanda Voermans, Aura Timen, Corine GeurtsvanKessel, Annemiek van der Eijk, Richard Molenkamp, Marion Koopmans, on behalf of the Dutch national COVID-19 response team. |
| EPI_ISL_415610 | hCoV-19/USA/WA-UW45/2020 | 690x | UW Virology Lab | UW Virology Lab | Pavitra Roychoudhury, Hong Xie, Keith Jerome, Alexander Greninger |
| EPI_ISL_415643 | hCoV-19/Georgia/Tb-468/2020 | 108x | R. G. Lugar Center for Public Health Research, National Center for Disease Control and Public Health (NCDC) of Georgia. | R. G. Lugar Center for Public Health Research, National Center for Disease Control and Public Health (NCDC) of Georgia. | Nato Kotaria, Marine Murtskhvaladze, Ann Machablishvili, Lela Sabadze, Mari Gavashelidze, Ana Papkiauri, Meri Pantsulaia, Gvantsa Brachveli, Tata Imnadze, Tamar Jashiashvili, Tea Tevdoradze, Ketevan Sidamonidze, Ekaterine Khmaladze, Ekaterine Zhghenti, Roena Sukhiashvili, Mariam Zakalashvili, Lela Urushadze, Magda Dgebuadze, Giorgi Tomashvili, Davit Tsaguria, Ekaterine Zangaladze, Nino Berishvili, Gvantsa Chanturia, Adam Kotorashvili, Maia Alkhazashvili, Irma Burjanadze, Anna Kasradze, Khatuna Zakhashvili, Paata Imnadze, |

| | | | | | |
|---|---|---|---|---|---|
| | | | | | Amiran Gamkrelidze. |
| EPI_ISL_416720 | hCoV-19/USA/WA-UW182/2020 | 175x | UW Virology Lab | UW Virology Lab | Pavitra Roychoudhury, Hong Xie, Keith Jerome, Alexander Greninger |
| EPI_ISL_417413 | hCoV-19/Turkey/6224-Ankara1034/2020 | 245x | Ministry of Health Turkey | Ministry of Health Turkey | Fatma Bayrakdar,Ayşe Başak Altaş,Yasemin Coşgun,Gülay Korukluoğlu,Selçuk Kılıç |
| EPI_ISL_418075 | hCoV-19/USA/WA-UW290/2020 | 151x | UW Virology Lab | UW Virology Lab | Pavitra Roychoudhury, Hong Xie, Keith Jerome, Alexander Greninger |
| EPI_ISL_418082 | hCoV-19/USA/WA-UW297/2020 | 291x | UW Virology Lab | UW Virology Lab | Pavitra Roychoudhury, Hong Xie, Keith Jerome, Alexander Greninger |
| EPI_ISL_418269 | hCoV-19/Vietnam/19-01S/2020 | 1897x | unknown | Microbiology and Immunology department | Cao,T.M., Nguyen,H.T., Pham,H.T.T., Vu,N.P.H., Dao,M.H., Huynh,L.T.K., Nguyen,L.T., Nguyen,N.T., Nguyen,T.T.N., Nguyen,A.H., Luong,Q.C., Nguyen,T.V., Tran,K.C., Pham,Q.D., Tran,T., Hoang,C.Q., Nguyen,T.T., Le,H.Q., Phung,T.M., Vo,T.N.A., Nguyen,S.N., Pham,D.T., Phan,L.T. and Nguyen,T.V. |
| EPI_ISL_419256 | hCoV-19/USA/VA-DCLS-0005/2020 | 1532x | Division of Consolidated Laboratory Services | Division of Consolidated Laboratory Services | Division of Consolidated Laboratory Services |
| EPI_ISL_419710 | hCoV-19/USA/VA-DCLS-0016/2020 | 4345x | Division of Consolidated Laboratory Services | Division of Consolidated Laboratory Services | Division of Consolidated Laboratory Services |
| EPI_ISL_419756 | hCoV-19/Australia/VIC38/2020 | 2983x | Victorian Infectious Diseases Reference Laboratory (VIDRL) | Victorian Infectious Diseases Reference Laboratory and Microbiological Diagnostic Unit Public Health Laboratory, Doherty Institute | Caly L., Seemann T., Sait, M., Schultz M., Druce J., Sherry, N. |
| EPI_ISL_419984 | hCoV-19/Australia/VIC293/2020 | 3158x | Victorian Infectious Diseases Reference Laboratory (VIDRL) | Victorian Infectious Diseases Reference Laboratory and Microbiological Diagnostic Unit Public | Caly L., Seemann T., Sait, M., Schultz M., Druce J., Sherry, N. |

| | | | | | |
|---|---|---|---|---|---|
| | | | | Health Laboratory, Doherty Institute | |
| EPI_ISL_420059 | hCoV-19/France/IDF3165/2020 | 13705x | Service de Biologie Médicale - BP 125 | National Reference Center for Viruses of Respiratory Infections, Institut Pasteur, Paris | Mélanie Albert, Marion Barbet, Sylvie Behillil, Méline Bizard, Angela Brisebarre, Flora Donati, Etienne Simon-Lorière, Vincent Enouf, Maud Vanpeene, Sylvie van der Werf, Christine Lambert |
| EPI_ISL_420181 | hCoV-19/England/SHEF-C0321/2020 | 7,000x, imputed from EPI_ISL_421275 and a large number of identities to others | Virology Department, Sheffield Teaching Hospitals NHS Foundation Trust | Department of Infection, Immunity and Cardiovascular Disease, The Florey Institute, The Medical School, University of Sheffield | Thushan de Silva, Matthew Parker, Adri Angyal, Rebecca Brown, Rachel Tucker, Paul Parsons, Luke Green, Danielle Groves, Alex Keeley, Dave Partridge, Matthew Wyles, Benjamin Lindsey, Mehmet Yavuz, Mohammad Raza, Cariad Evans |
| EPI_ISL_420182 | hCoV-19/England/SHEF-C029D/2020 | Coverage imputed by identity to EPI_ISL_420195 | Virology Department, Sheffield Teaching Hospitals NHS Foundation Trust | Department of Infection, Immunity and Cardiovascular Disease, The Florey Institute, The Medical School, University of Sheffield | Thushan de Silva, Matthew Parker, Adri Angyal, Rebecca Brown, Rachel Tucker, Paul Parsons, Luke Green, Danielle Groves, Alex Keeley, Dave Partridge, Matthew Wyles, Benjamin Lindsey, Mehmet Yavuz, Mohammad Raza, Cariad Evans |
| EPI_ISL_420241 | hCoV-19/England/SHEF-C0637/2020 | 2090x, imputed from EPI_ISL_417084 | Virology Department, Sheffield Teaching Hospitals NHS Foundation Trust | Department of Infection, Immunity and Cardiovascular Disease, The Florey Institute, The Medical School, University of Sheffield | Thushan de Silva, Matthew Parker, Adri Angyal, Rebecca Brown, Rachel Tucker, Paul Parsons, Luke Green, Danielle Groves, Alex Keeley, Dave Partridge, Matthew Wyles, Benjamin Lindsey, Mehmet Yavuz, Mohammad Raza, Cariad Evans |
| EPI_ISL_420306 | hCoV-19/USA/AK-PHL015/2020 | 1666x, imputed from EPI_ISL_419565 | Alaska State Virology Laboratory | Alaska State Virology Laboratory | Chen, J. |
| EPI_ISL_420422 | hCoV-19/Belgium/JJC-0325165/2020 | 13,550x | KU Leuven, Clinical and Epidemiological Virology | KU Leuven, Clinical and Epidemiological Virology | Joan Marti-Carreras, Bert Vanmechelen, Tony Wawina, Piet Maes |
| EPI_ISL_420510 | hCoV-19/England/201320 | coverage unknown, imputing by identity | Respiratory Virus Unit, Microbiology Services Colindale, | Respiratory Virus Unit, Microbiology | Monica Galiano, Shahjahan Miah, Angie Lackenby, Omolola Akinbami, Tiina Talts, Leena |

| | | | | | |
|---|---|---|---|---|---|
| | 74102/2020 | to EPI_ISL_420509 | Public Health England | Services Colindale, Public Health England | Bhaw, Richard Myers, Steven Platt, Kirstin Edwards, Jonathan Hubb, Joanna Ellis, Maria Zambon |
| EPI_ISL_420579 | hCoV-19/USA/NY-NYUMC64/2020 | 298x | NYU Langone Health | Departments of Pathology and Medicine, New York University School of Medicine | Maria Aguero-Rosenfeld, Brendan Belovarac, Margaret Black, Ludovic Boytard, John Cadley, Paolo Cotzia, John Chen, Dacia Dimartino, Xiaojun Feng, Tatyana Gindin, Adriana Heguy, Megan Hogan, Emily Huang, George Jour, Andrew Lytle, Christian Marier, Matthew T. Maurano, Mark J. Mulligan, Peter Meyn, Iman Osman, Jared Pinnell, Sitharam Ramaswami, Amy Rapkiewicz, Marie Samanovic-Golden, Antonio Serrano, Guomiao Shen, Matija Snuderl, Theodore Vougiouklakis, Nick Vulpescu, Gael Westby, Paul Zappile, Yutong Zhang |
| EPI_ISL_420610 | hCoV-19/France/ARA12558/2020 | 3,784x | Institut des Agents Infectieux (IAI), Hospices Civils de Lyon | CNR Virus des Infections Respiratoires - France SUD | Antonin Bal, Gregory Destras, Gwendolyne Burfin, Solenne Brun, Carine Moustaud, Raphaelle Lamy, Alexandre Gaymard, Maude Bouscambert-Duchamp, Florence Morfin-Sherpa, Martine Valette, Bruno Lina, Laurence Josset |
| EPI_ISL_421005 | hCoV-19/Wales/PHWC-25560/2020 | 3,213x, imputing from EPI_ISL_424455 and multiple identities | Wales Specialist Virology Centre | Public Health Wales Microbiology Cardiff | Catherine Moore, Joanne Watkins, Sally Corden, Malorie Perry, Simon Cottrell Sara Rey, Matt Bull, Tom Connor |
| EPI_ISL_421312 | hCoV-19/USA/WI-54/2020 | 400x | University of Wisconsin-Madison AIDS Vaccine Research Laboratories | University of Wisconsin-Madison AIDS Vaccine Research Laboratories | Gage Moreno, Katarina Braun |
| EPI_ISL_421380 | hCoV-19/USA/NY-PV08417/2020 | 3017x, imputed from EPI_ISL_420012 and multiple identities | MSHS Clinical Microbiology Laboratories | MSHS Pathogen Surveillance Program | Ana S. Gonzalez-Reiche, Mitchell Sullivan, Ajay Obla, Gopi Patel, Emilia Sordillo, Melissa Gitman, Alberto Paniz-mondolfi, Matthew Hernandez, Shelcie Fabre, Jose Polanco, Zenab Khan, Bremy Albuquerque, Jayeeta Dutta, Juan |

| | | | | | |
|---|---|---|---|---|---|
| | | | | | Soto, Shwetha Sridhar Hara, Ying-Chih Wang, Melissa Smith, Robert Sebra, Lisa Miorin, Wen-chun Liu, Randy Albrecht, Judith Aberg, Florian Krammer, Adolfo Garcia-Sarstre, Viviana Simon, Harm van Bakel |
| EPI_ISL_421506 | hCoV-19/France/IDF3236/2020 | 99.69% covered 241x | Service de Biologie Médicale - BP 125 | National Reference Center for Viruses of Respiratory Infections, Institut Pasteur, Paris | Mélanie Albert, Marion Barbet, Sylvie Behillil, Méline Bizard, Angela Brisebarre, Flora Donati, Etienne Simon-Lorière, Vincent Enouf, Maud Vanpeene, Sylvie van der Werf, Christine Lambert |
| EPI_ISL_421515 | hCoV-19/Spain/Valencia40/2020 | 434x | Servicio de Microbiologia. Hospital Clinico Universitario de Valencia | Sequencing and Bioinformatics Service and Molecular Epidemiology Research Group. FISABIO-Public Health | Giuseppe D'Auria, Llúcia Martínez-Priego, Maria Alma Bracho, Griselda De Marco, Beatriz Beamud, Lidia Ruiz Roldan, Marta Pla Diaz, Neris Garcia-Gonzalez, Loreto Ferrús Abad, Inma Galán Vendrell, Paula Ruiz-Hueso, Mariana Reyes-Prieto, Vicente Soriano Chirona, David Navarro, Fernando Gonzalez-Candelas |
| EPI_ISL_421763 | hCoV-19/Luxembourg/LNS0074148/2020 | 4,808x | Laboratoire National de Sante, Microbiology, Virology | Laboratoire National de Sante, Microbiology, Epidemiology and Microbial Genomics | Anke Wienecke-Baldacchino, Ardashel Latsuzbaia, Jessica Tapp, Catherine Ragimbeau, Guillaume Fournier, Tamir Abdelrahman, Trung Nguyen Nguyen, Joel Mossong |
| EPI_ISL_422184 | hCoV-19/Wales/PHWC-26875/2020 | Coverage imputed from identity to EPI_ISL_422052 | Wales Specialist Virology Centre | Public Health Wales Microbiology Cardiff | Catherine Moore, Johnathan Evans, Malorie Perry, Simon Cottrell, Alec Birchley, Alexander Adams, Amy Gaskin, Bree Gatica-Wilcox, Jason Coombes, Lauren Gilbert, Lee Graham, Nicole Pacchiarini, Sara Kumziene-Summerhayes, Sarah Taylor, Sophie Jones, Sara Rey, Matthew Bull, Joanne Watkins, Sally Corden, Tom Connor |
| EPI_ISL_422860 | hCoV-19/Netherlands/NoordBrabant_101/2020 | 30x | Dutch COVID-19 response team | Erasmus Medical Center | Bas Oude Munnink, David Nieuwenhuijse, Reina Sikkema, Claudia Schapendonk, Irina Chestakova, Anne van der Linden, Theo Bestebroer, Stefan van Nieuwkoop, Mark Pronk, Pascal Lexmond, Corien Swaan, Manon Haverkate, Madelief Mollers, Mart Stein, Sandra |

| | | | | | |
|---|---|---|---|---|---|
| | | | | | Kengne Kamga Mobou, Jeroen van Kampen, Jolanda Voermans, Aura Timen, Corine GeurtsvanKessel, Annemiek van der Eijk, Richard Molenkamp, Marion Koopmans, on behalf of the Dutch national COVID-19 response team. |
| EPI_ISL_422919 | hCoV-19/Netherlands/ZuidHolland_48/2020 | 30x | Dutch COVID-19 response team | Erasmus Medical Center | Bas Oude Munnink, David Nieuwenhuijse, Reina Sikkema, Claudia Schapendonk, Irina Chestakova, Anne van der Linden, Theo Bestebroer, Stefan van Nieuwkoop, Mark Pronk, Pascal Lexmond, Corien Swaan, Manon Haverkate, Madelief Mollers, Mart Stein, Sandra Kengne Kamga Mobou, Jeroen van Kampen, Jolanda Voermans, Aura Timen, Corine GeurtsvanKessel, Annemiek van der Eijk, Richard Molenkamp, Marion Koopmans, on behalf of the Dutch national COVID-19 response team. |
| EPI_ISL_423007 | hCoV-19/USA/WA-UW-1448/2020 | 7,813x, imputed from EPI_ISL_424606 | UW Virology Lab | UW Virology Lab | Pavitra Roychoudhury, Hong Xie, Keith Jerome, Alexander Greninger |
| EPI_ISL_423288 | hCoV-19/England/20138017804/2020 | Coverage imputed by identity to EPI_ISL_421979, EPI_ISL_420529, EPI_ISL_420702, EPI_ISL_416907 | Respiratory Virus Unit, Microbiology Services Colindale, Public Health England | Respiratory Virus Unit, Microbiology Services Colindale, Public Health England | Monica Galiano, Shahjahan Miah, Angie Lackenby, Omolola Akinbami, Tiina Talts, Leena Bhaw, Richard Myers, Steven Platt, Kirstin Edwards, Jonathan Hubb, Joanna Ellis, Maria Zambon |
| EPI_ISL_423642 | hCoV-19/England/20130062404/2020 | Coverage imputed by identity to EPI_ISL_424129 | Respiratory Virus Unit, Microbiology Services Colindale, Public Health England | Respiratory Virus Unit, Microbiology Services Colindale, Public Health England | Monica Galiano, Shahjahan Miah, Angie Lackenby, Omolola Akinbami, Tiina Talts, Leena Bhaw, Richard Myers, Steven Platt, Kirstin Edwards, Jonathan Hubb, Joanna Ellis, Maria Zambon |
| EPI_ISL_423725 | hCoV-19/England/20132005904/2020 | Coverage imputed by identity to EPI_ISL_423267 | Respiratory Virus Unit, Microbiology Services Colindale, Public Health England | Respiratory Virus Unit, Microbiology Services Colindale, | Monica Galiano, Shahjahan Miah, Angie Lackenby, Omolola Akinbami, Tiina Talts, Leena Bhaw, Richard Myers, Steven Platt, Kirstin |

| | | | | | |
|---|---|---|---|---|---|
| | | | | Public Health England | Edwards, Jonathan Hubb, Joanna Ellis, Maria Zambon |
| EPI_ISL_423772 | hCoV-19/England/20132032604/2020 | Coverage imputed by identity to EPI_ISL_421903 | Respiratory Virus Unit, Microbiology Services Colindale, Public Health England | Respiratory Virus Unit, Microbiology Services Colindale, Public Health England | Monica Galiano, Shahjahan Miah, Angie Lackenby, Omolola Akinbami, Tiina Talts, Leena Bhaw, Richard Myers, Steven Platt, Kirstin Edwards, Jonathan Hubb, Joanna Ellis, Maria Zambon |
| EPI_ISL_424470 | hCoV-19/Iceland/446/2020 | 2,385x | The National University Hospital of Iceland | deCODE genetics | Daniel F Gudbjartsson; Agnar Helgason; Hakon Jonsson; Olafur T Magnusson; Pall Melsted; Gudmundur L Norddahl; Jona Saemundsdottir; Asgeir Sigurdsson; Patrick Sulem; Arna B Agustsdottir; Berglind Eiriksdottir; Run Fridriksdottir; Elisabet E Gardarsdottir; Gudmundur Georgsson; Olafia S Gretarsdottir; Kjartan R Gudmundsson; Thora R Gunnarsdottir; Arnaldur Gylfason; Hilma Holm; Brynjar O Jensson; Aslaug Jonasdottir; Kamilla S Josefsdottir; Thordur Kristjansson; Droplaug N Magnusdottir; Louise le Roux; Gudrun Sigmundsdottir; Gardar Sveinbjornsson; Kristin E Sveinsdottir; Maney Sveinsdottir; Emil A Thorarensen; Bjarni Thorbjornsson; Gisli Masson; Ingileif Jonsdottir; Alma Moller; Thorolfur Gudnason; Karl G Kristinsson; Unnur Thorsteinsdottir; Kari Stefansson |
| EPI_ISL_425132 | hCoV-19/Germany/NRW-48/2020 | 1,000x | Center of Medical Microbiology, Virology, and Hospital Hygiene, University of Duesseldorf | Center of Medical Microbiology, Virology, and Hospital Hygiene, University of Duesseldorf | Ortwin Adams, Marcel Andree, Alexander Dilthey, Torsten Feldt, Sandra Hauka, Torsten Houwaart, Bjorn-Erik Jensen, Detlef Kindgen-Milles, Malte Kohns Vasconcelos, Klaus Pfeffer, Tina Senff, Daniel Strelow, Jorg Timm, Andreas Walker, Tobias Wienemann |
| EPI_ISL_425235 | hCoV-19/England/CAMB-7378B/2020 | 3,446.95x | Department of Pathology, University of Cambridge | COVID-19 Genomics UK (COG-UK) Consortium | Luke W Meredith, M. Estee Torok , Myra Hosmillo, William L. Hamilton, Martin D. Curran, Theresa Feltwell, Anna Yakovleva, |

| | | | | | |
|---|---|---|---|---|---|
| | | | | | Charlotte J. Houldcroft, Aminu S. Jahun, Sarah L. Caddy, Ian Goodfellow |
| EPI_ISL_425242 | hCoV-19/England/CAMB-737F4/2020 | 5,515.73x | Department of Pathology, University of Cambridge | COVID-19 Genomics UK (COG-UK) Consortium | Luke W Meredith, M. Estee Torok , Myra Hosmillo, William L. Hamilton, Martin D. Curran, Theresa Feltwell, Anna Yakovleva, Charlotte J. Houldcroft, Aminu S. Jahun, Sarah L. Caddy, Ian Goodfellow |
| EPI_ISL_425284 | hCoV-19/England/CAMB-74395/2020 | 3,693.48x | Department of Pathology, University of Cambridge | COVID-19 Genomics UK (COG-UK) Consortium | Luke W Meredith, M. Estee Torok , Myra Hosmillo, William L. Hamilton, Martin D. Curran, Theresa Feltwell, Anna Yakovleva, Charlotte J. Houldcroft, Aminu S. Jahun, Sarah L. Caddy, Ian Goodfellow |
| EPI_ISL_425319 | hCoV-19/England/CAMB-74650/2020 | 1,814.79x | Department of Pathology, University of Cambridge | COVID-19 Genomics UK (COG-UK) Consortium | Luke W Meredith, M. Estee Torok , Myra Hosmillo, William L. Hamilton, Martin D. Curran, Theresa Feltwell, Anna Yakovleva, Charlotte J. Houldcroft, Aminu S. Jahun, Sarah L. Caddy, Ian Goodfellow |
| EPI_ISL_425342 | hCoV-19/England/CAMB-74A09/2020 | 1,554.04x | Department of Pathology, University of Cambridge | COVID-19 Genomics UK (COG-UK) Consortium | Luke W Meredith, M. Estee Torok , Myra Hosmillo, William L. Hamilton, Martin D. Curran, Theresa Feltwell, Anna Yakovleva, Charlotte J. Houldcroft, Aminu S. Jahun, Sarah L. Caddy, Ian Goodfellow |
| EPI_ISL_425498 | hCoV-19/England/NOTT-10E0F2/2020 | 8,508.72x | Queens Medical Centre, Clinical Microbiology Department / DeepSeq Nottingham | COVID-19 Genomics UK (COG-UK) Consortium | Gemma Clark, Wendy Smith, Manjinder Khakh, Hannah Howson-Wells, Jonathan Ball, Patrick McClure, Joseph Chappell, Theocharis Tsoleridis, Nadine Holmes, Matthew Carlisle, Christopher Moore, Fei Sang, Johnny Debebe, Victoria Wright, Matthew Loose |
| EPI_ISL_425643 | hCoV-19/England/NOTT-10EB3D/2020 | 5,122.9x | Queens Medical Centre, Clinical Microbiology Department / DeepSeq Nottingham | COVID-19 Genomics UK (COG-UK) Consortium | Gemma Clark, Wendy Smith, Manjinder Khakh, Hannah Howson-Wells, Jonathan Ball, Patrick McClure, Joseph Chappell, Theocharis Tsoleridis, Nadine Holmes, Matthew Carlisle, Christopher Moore, Fei Sang, Johnny Debebe, Victoria Wright, Matthew Loose |
| EPI_ISL_425655 | hCoV- | 3,671.82x | West of Scotland Specialist | COVID-19 Genomics | Ana da Silva Filipe, Kathy Smollett, Stephen |

| | | | | | |
|---|---|---|---|---|---|
| | 19/Scotland/CVR11/2020 | | Virology Centre, NHSGGC / MRC-University of Glasgow Centre for Virus Research | UK (COG-UK) Consortium | Carmichael, Natasha Johnson, Daniel Mair, Lily Tong, Jenna Nichols; Sarah McDonald; Richard Orton, Joseph Hughes, Sreenu Vattipally, David L Robertson; Kathy Li, Natasha Jesudason, Rajiv Shah, James Shepherd, Antonia Ho, Emma Thomson; Alasdair MacLean, Rory Gunson. |
| EPI_ISL_425839 | hCoV-19/Scotland/EDB033/2020 | 420.12x | Virology Department, Royal Infirmary of Edinburgh, NHS Lothian / School of Biological Sciences, University of Edinburgh / Institute of Genetics and Molecular Medicine, University of Edinburgh | COVID-19 Genomics UK (COG-UK) Consortium | McHugh M, Dewar R, Rooke S, Gallagher M, Balcaza C, O'Toole A, Hill V, McCrone JT, Colquhoun R, Yu X, Jackson B, Scher E, Rambaut A, Williams TC, Templeton K |
| EPI_ISL_425886 | hCoV-19/Scotland/EDB096/2020 | 423.97x | Virology Department, Royal Infirmary of Edinburgh, NHS Lothian / School of Biological Sciences, University of Edinburgh / Institute of Genetics and Molecular Medicine, University of Edinburgh | COVID-19 Genomics UK (COG-UK) Consortium | McHugh M, Dewar R, Rooke S, Gallagher M, Balcaza C, O'Toole A, Hill V, McCrone JT, Colquhoun R, Yu X, Jackson B, Scher E, Rambaut A, Williams TC, Templeton K |
| EPI_ISL_426028 | hCoV-19/USA/NY-Wadsworth-10690-01/2020 | 9,883x | Wadsworth Center, New York State Department.of Health | Wadsworth Center, New York State Department.of Health | Kirsten St. George, Daryl M. Lamson, Sara Griesemer, Jonathan Plitnick, Navjot Singh, Matthew D. Shudt, Erica Lasek-Nesselquist |
| EPI_ISL_426097 | hCoV-19/USA/WA-UW-2142/2020 | Coverage imputed by identity to EPI_ISL_424166 | UW Virology Lab | UW Virology Lab | Pavitra Roychoudhury, Hong Xie, Keith Jerome, Alexander Greninger |

**Supplementary Table A2: High *BC* residues shown for each chain (A and B)**

| Accession number | High *BC* residues (top 5% ECD) |
|---|---|
| EPI_ISL_413021 | **17, 128,** 111, 112, 14, 7, 3, 140, 115, 11, 163, 129, 148, 138, 290 |
| | **128, 17,** 18, 115, 146, 126, 11, 14, 39, 148, 13, 147, 136, 140, 127 |
| EPI_ISL_415503 | **17, 128,** 115, 11, 13, 148, 18, 14, 116, 6, 147, 150, 111, 146, 112 |
| | **128, 17,** 115, 39, 112, 290, 18, 147, 14, 111, 148, 13, 6, 129, 11 |
| EPI_ISL_415610 | **17, 128,** 14, 111, 115, 112, 3, 147, 39, 18, 126, 6, 11, 296, 146 |
| | **17, 128,** 11, 13, 112, 115, 111, 146, 39, 18, 148, 14, 299, 147, 296 |
| EPI_ISL_415643 | **17, 128,** 111, 115, 112, 3, 14, 18, 141, 39, 11, 148, 299, 292, 129 |
| | 115, **17,** 111, 139, **128,** 112, 7, 39, 18, 13, 127, 11, 148, 14, 4 |
| EPI_ISL_416720 | **17,** 115, 14, 112, 39, 6, 146, 127, 111, 147, 140, 163, 18, **128,** 11 |
| | 6, **17,** 5, 146, 111, 112, **128,** 292, 115, 39, 14, 127, 282, 140, 138 |
| EPI_ISL_417413 | **17, 128,** 112, 3, 140, 111, 116, 14, 129, 13, 18, 115, 148, 6, 11 |
| | **17,** 115, **128,** 6, 140, 139, 2, 146, 112, 39, 111, 147, 291, 18, 163 |
| EPI_ISL_418075 | **17, 128,** 111, 139, 112, 3, 18, 115, 11, 14, 163, 140, 292, 13, 148 |
| | **17, 128,** 115, 39, 18, 111, 112, 139, 148, 14, 146, 140, 3, 13, 124 |
| EPI_ISL_418082 | **17,** 111, 112, 140, 14, 3, 139, 11, **128,** 13, 115, 292, 148, 116, 18 |
| | **17, 128,** 115, 14, 129, 111, 146, 148, 147, 286, 112, 39, 11, 18, 140 |
| EPI_ISL_418269 | **128, 17,** 39, 290, 13, 11, 147, 126, 18, 112, 2, 148, 146, 14, 115 |
| | **17, 128,** 115, 14, 39, 147, 299, 18, 140, 111, 112, 6, 284, 296, 129 |
| EPI_ISL_419256 | **17, 128,** 115, 18, 6, 14, 112, 127, 290, 147, 286, 13, 2, 148, 111 |
| | 111, 115, **17,** 112, 6, 140, **128,** 139, 2, 126, 146, 5, 116, 290, 292 |
| EPI_ISL_419710 | **128, 17,** 115, 163, 11, 6, 114, 14, 18, 3, 126, 290, 13, 111, 112 |
| | **17,** 111, 14, 18, 139, 112, **128,** 13, 115, 11, 146, 292, 6, 284, 5 |
| EPI_ISL_419756 | **17,** 146, 11, 115, 139, 14, 127, 39, 130, 18, 140, 3, 6, 13, 136 |
| | 111, **17,** 112, 115, 14, 3, **128,** 11, 6, 18, 148, 146, 39, 13, 140 |
| EPI_ISL_419984 | **17,** 115, **128,** 292, 111, 11, 112, 13, 14, 162, 116, 150, 299, 296, 126 |
| | **17,** 282, 4, 18, **128,** 115, 14, 13, 148, 11, 139, 39, 6, 111, 284 |
| EPI_ISL_420059 | **17, 128,** 290, 111, 14, 112, 115, 18, 148, 126, 147, 139, 7, 146, 13 |
| | **17,** 115, 111, 112, 18, 13, **128,** 11, 3, 148, 39, 126, 146, 122, 140 |
| EPI_ISL_420181 | **17,** 112, 111, **128,** 140, 14, 39, 18, 115, 146, 139, 11, 147, 148, 126 |
| | **128,** 115, **17,** 147, 146, 299, 112, 2, 290, 14, 139, 140, 111, 296, 39 |
| EPI_ISL_420182 | **128, 17,** 290, 6, 115, 147, 140, 39, 14, 146, 129, 126, 114, 18, 111 |
| | **17,** 112, 115, 111, **128,** 39, 14, 18, 3, 147, 292, 146, 148, 141, 116 |
| EPI_ISL_420241 | **128, 17,** 6, 11, 111, 5, 126, 112, 115, 18, 13, 282, 286, 290, 148 |
| | **17,** 39, 115, 14, **128,** 18, 116, 139, 146, 140, 144, 127, 6, 112, 147 |
| EPI_ISL_420306 | **17,** 14, 111, 112, 115, 141, 39, 18, **128,** 3, 146, 147, 13, 11, 116 |
| | **17, 128,** 11, 111, 13, 6, 148, 18, 115, 146, 147, 112, 14, 288, 296 |
| EPI_ISL_420422 | **17,** 139, 111, 115, 112, 14, 18, 140, 6, **128,** 163, 127, 292, 214, 7 |
| | **128, 17,** 111, 112, 11, 39, 115, 14, 18, 13, 3, 129, 148, 139, 299 |
| EPI_ISL_420510 | **128,** 111, 112, 115, **17,** 147, 13, 116, 148, 11, 291, 39, 18, 146, 14 |
| | **128, 17,** 290, 115, 6, 147, 14, 39, 111, 146, 112, 18, 116, 286, 127 |
| EPI_ISL_420579 | **17,** 112, 111, 14, **128,** 148, 115, 146, 39, 147, 150, 13, 18, 139, 116 |
| | **128, 17,** 115, 111, 290, 3, 18, 126, 112, 11, 148, 150, 13, 139, 14 |
| EPI_ISL_420610 | **17,** 299, 296, 115, 39, 11, 146, 14, 147, 13, 111, 18, 148, 127, 112 |
| | **128, 17,** 290, 140, 39, 111, 115, 14, 3, 11, 148, 112, 126, 18, 13 |
| EPI_ISL_421005 | 112, **17,** 111, 141, 115, 292, 14, 39, 147, 146, 214, 140, 13, 18, 128 |
| | 111, 112, **17,** 292, 115, **128,** 163, 296, 148, 299, 150, 18, 13, 162, 290 |
| EPI_ISL_421312 | **17, 128,** 115, 290, 6, 18, 147, 112, 14, 111, 13, 11, 148, 146, 296 |
| | **17,** 111, 115, 140, 112, 18, 14, 163, **128,** 141, 126, 39, 146, 7, 6 |
| EPI_ISL_421380 | **17, 128,** 127, 111, 139, 290, 7, 14, 11, 146, 13, 18, 112, 115, 148 |
| | 39, 112, **17,** 3, 115, 148, 147, 111, **128,** 13, 14, 292, 11, 18, 146 |

| | |
|---|---|
| EPI_ISL_421506 | **17,** 112, 111, 115, 14, 18, 139, 162, 13, 148, 126, 11, 39, 292, 285 |
| | **17,** 111, 115, 112, 7, 14, 18, **128,** 146, 139, 147, 127, 148, 39, 214 |
| EPI_ISL_421515 | **17, 128,** 6, 115, 14, 111, 112, 147, 292, 290, 18, 11, 39, 3, 296 |
| | **17,** 111, 112, 14, 115, 139, 116, 146, 18, **128,** 147, 292, 39, 163, 140 |
| EPI_ISL_421763 | **17, 128,** 111, 115, 112, 14, 290, 11, 18, 39, 147, 116, 13, 148, 6 |
| | **17,** 112, 111, **128,** 115, 139, 148, 14, 18, 146, 39, 140, 290, 11, 129 |
| EPI_ISL_422184 | **17,** 112, 115, 111, 14, 18, 39, **128,** 140, 6, 292, 299, 163, 147, 148 |
| | **128, 17,** 115, 11, 14, 127, 18, 2, 112, 7, 140, 147, 290, 39, 113 |
| EPI_ISL_422860 | **17,** 111, 112, 115, 18, 299, 292, 140, 163, 126, 39, 5, 139, 14, 146 |
| | **17,** 115, **128,** 141, 39, 112, 14, 116, 140, 129, 147, 148, 111, 290, 2 |
| EPI_ISL_422919 | **17,** 146, 112, 115, 111, **128,** 14, 39, 11, 299, 296, 148, 290, 18, 13 |
| | **17, 128,** 115, 14, 39, 111, 112, 3, 141, 18, 6, 11, 290, 148, 147 |
| EPI_ISL_423007 | **17, 128,** 6, 286, 140, 112, 13, 111, 115, 139, 129, 11, 127, 5, 18 |
| | **17,** 144, 39, 115, 141, 6, 14, 111, **128,** 214, 292, 286, 112, 18, 127 |
| EPI_ISL_423288 | **17,** 39, **128,** 129, 6, 14, 112, 3, 11, 146, 13, 111, 148, 18, 115 |
| | **128, 17,** 290, 39, 146, 111, 11, 139, 13, 148, 14, 112, 18, 140, 126 |
| EPI_ISL_423642 | **17,** 111, 112, 115, 139, 140, 14, 7, **128,** 18, 6, 292, 126, 5, 3 |
| | **17,** 127, 115, 111, **128,** 112, 18, 14, 11, 148, 13, 6, 139, 150, 146 |
| EPI_ISL_423725 | **17,** 112, 111, 5, 14, 148, 115, **128,** 11, 286, 4, 13, 18, 141, 3 |
| | **17, 128,** 14, 112, 111, 286, 39, 18, 115, 299, 13, 129, 11, 6, 296 |
| EPI_ISL_423772 | **17, 128,** 111, 112, 18, 115, 147, 290, 11, 14, 116, 39, 148, 13, 129 |
| | **17, 128,** 115, 5, 14, 18, 148, 11, 6, 124, 163, 291, 290, 111, 146 |
| EPI_ISL_424470 | **17,** 115, **128,** 147, 3, 111, 112, 139, 18, 14, 129, 39, 148, 288, 11 |
| | **17,** 139, **128,** 290, 111, 14, 18, 6, 115, 11, 112, 5, 39, 13, 148 |
| EPI_ISL_425132 | 139, **17,** 11, 140, 115, 112, 146, 111, **128,** 7, 147, 13, 282, 116, 14 |
| | **17,** 112, 111, 39, **128,** 115, 14, 18, 13, 3, 148, 299, 146, 139, 11 |
| EPI_ISL_425235 | **17, 128,** 14, 115, 146, 139, 112, 111, 39, 18, 3, 140, 148, 13, 11 |
| | **128,** 11, **17,** 39, 111, 3, 13, 148, 112, 162, 140, 139, 147, 163, 299 |
| EPI_ISL_425242 | **17,** 11, **128,** 139, 39, 112, 111, 115, 14, 13, 148, 18, 124, 296, 7 |
| | **17,** 14, 112, 115, 111, 39, **128,** 18, 3, 299, 7, 127, 129, 147, 296 |
| EPI_ISL_425284 | **17,** 299, 115, 111, 39, 14, 296, 116, 6, 112, 146, 126, 18, 292, 128 |
| | **17, 128,** 6, 124, 14, 115, 141, 290, 11, 18, 13, 39, 145, 2, 291 |
| EPI_ISL_425319 | **17, 128,** 7, 6, 115, 111, 14, 112, 18, 13, 139, 126, 148, 286, 11 |
| | 112, **17,** 111, 126, **128,** 14, 7, 115, 139, 140, 148, 3, 299, 18, 146 |
| EPI_ISL_425342 | **17,** 139, **128,** 111, 112, 115, 5, 14, 290, 6, 3, 18, 11, 13, 148 |
| | **128, 17,** 115, 11, 146, 18, 111, 112, 147, 148, 3, 13, 14, 39, 6 |
| EPI_ISL_425498 | **17, 128,** 115, 111, 112, 14, 18, 6, 139, 13, 148, 147, 11, 140, 3 |
| | **128, 17,** 127, 115, 147, 6, 126, 116, 14, 139, 129, 114, 290, 140, 296 |
| EPI_ISL_425643 | 6, **128, 17,** 290, 11, 111, 112, 39, 5, 13, 18, 148, 127, 115, 14 |
| | **17,** 115, **128,** 14, 299, 18, 296, 111, 112, 2, 13, 292, 3, 147, 116 |
| EPI_ISL_425655 | **17, 128,** 290, 115, 112, 111, 14, 39, 18, 148, 11, 140, 288, 7, 13 |
| | **17,** 112, 115, 14, 111, **128,** 18, 13, 39, 292, 148, 11, 147, 3, 7 |
| EPI_ISL_425839 | **128, 17,** 39, 11, 18, 148, 115, 13, 111, 147, 112, 14, 2, 146, 126 |
| | **17,** 111, 115, 139, 112, 14, 18, 39, **128,** 126, 3, 140, 213, 13, 292 |
| EPI_ISL_425886 | **17,** 111, 112, 115, 14, 292, 39, 146, 172, 299, 296, 148, 163, 18, 6 |
| | **17,** 139, 283, 39, 11, 146, 140, 284, 163, 14, 148, 18, 7, 112, 126 |
| EPI_ISL_426028 | **17,** 163, 127, 14, 296, 112, 11, **128,** 18, 299, 3, 13, 111, 7, 140 |
| | **17,** 139, 115, 7, 127, 111, 18, **128,** 112, 140, 288, 13, 11, 148, 290 |
| EPI_ISL_426097 | **17,** 6, **128,** 115, 14, 39, 5, 112, 3, 111, 18, 11, 116, 286, 129 |
| | **17,** 139, **128,** 115, 112, 14, 111, 39, 286, 5, 140, 11, 148, 13, 18 |
| Reference | **17,** 139, 112, 111, 18, 115, 14, 11, 7, **128,** 13, 148, 127, 290, 140 |
| | **17,** 115, **128,** 14, 148, 146, 11, 6, 18, 129, 111, 13, 126, 3, 140 |
